# Understanding Spatial Patterns of Human-Tiger and Human-Leopard Conflict across South Asia using a Multiscale Regional Approach

**DOI:** 10.64898/2026.09.26.754692

**Authors:** Anish Banerjee

## Abstract

Over the past two decades, significant population recoveries of tigers and leopards have driven a substantial rise in human-carnivore conflict across South Asia. Sustaining the long-term conservation of both species demands effective, adaptive conflict management. However, existing research is largely confined to isolated landscapes or protected areas, hindering the identification of spatial conflict drivers across broader, heterogeneous regions. To address this gap, this study compiled 4,457 human-leopard conflict (HLC) and 2,563 human-tiger conflict (HTC) records spanning 2000-2025 across Bangladesh, Bhutan, India, and Nepal. We evaluated conflict locations against 33 landscape, anthropogenic, climatic, topographic, and hydrological predictors across three spatial scales using binomial generalized linear regression models and three alternative background sampling methods. Model performance peaked at the largest spatial scale for both species and remained consistent across sampling strategies. HTC correlated positively with forest fragmentation and proximity to human settlements, but decreased with higher overall anthropogenic disturbance. Conversely, HLC concentrated in highly productive, high-precipitation, human-modified landscapes. These contrasting spatial patterns highlight distinct ecological responses to habitat modification, carrying unique conservation implications. Ultimately, this study underscores the necessity of integrating carnivore abundance, wild prey densities, and socioeconomic factors to achieve a holistic understanding of conflict dynamics.

## 1. Introduction

Tiger and leopard populations have recovered substantially across South Asia in recent decades, despite significant socioeconomic pressures (Jhala et al., 2021; Pandey, Apan and Maraseni, 2026; Qureshi et al., 2024). However, this conservation success has intensified human-carnivore conflict, resulting in substantial costs, including human casualties, livestock losses, and associated livelihood losses (Nature Conservation Division, 2024). Specifically, between 2000 and 2005, there were 459 reported human fatalities due to tiger attacks in India (Lok Sabha, 2026). Similar patterns have been observed in neighboring countries such as Bhutan, Bangladesh, and Nepal (Dhar and Mondal, 2023; Rostro-García *et al*., 2016; Shrestha *et al*., 2026). These negative interactions can reduce local tolerance towards large carnivores, often resulting in retaliatory killing (Pandey, Apan and Maraseni, 2026). Effective adaptive management is therefore required for the sustained long-term conservation of both species and requires a comprehensive understanding of the spatial determinants of human-felid conflict (Kuiper *et al*., 2024). Landscape structure, anthropogenic disturbances, bioclimatic, hydrological, and topographic conditions all significantly influence conflict distribution.

Landscape structure and land-use change influence human-carnivore conflict patterns by altering carnivore distribution and movement, often increasing spatial overlap with people and livestock (Dutta *et al*., 2020). For example, forest fragmentation has been associated with increased livestock depredation, particularly in forest patches interspersed with agricultural land, grazing areas, and settlements (Acharya *et al*., 2017; Kshettry, Vaidyanathan and Athreya, 2018; Sharma *et al*., 2020). Similarly, human-carnivore conflict has often increased after substantial habitat loss and degradation, as reduced wild prey densities increase reliance on domestic prey (Inskip and Zimmermann, 2009; Bhattacharya *et al*., 2025). However, greater canopy and forest cover has been associated with increased conflict for both species (Chatterjee *et al*., 2022; Soh et al., 2014a). Both species rely on more structurally complex vegetation for concealment and hunting opportunities, while sufficient forest habitat supports large-felid persistence and movement through human-modified landscapes (Puri *et al*., 2020; Yumnam *et al*., 2014).

Anthropogenic pressure further modifies these spatial patterns, with tigers and leopards responding differently. Human-leopard conflict has been associated with greater proximity to settlements and increased built-up area (Akrim *et al*., 2023; Lamichhane *et al*., 2023; Naha *et al*., 2020). Leopards, due to their behavioral and ecological plasticity, have persisted in agricultural and other human-modified landscapes by exploiting anthropogenic food resources, particularly free-ranging dogs and livestock (Athreya *et al*., 2014; Odden *et al*., 2014). Moreover, human-leopard conflict was higher near roads across the Himalayan landscapes, potentially reflecting increased access to leopard habitat by people and livestock (Sharma *et al*., 2020; Silwal *et al*., 2024). In contrast, road proximity has shown more variable relationships with human-tiger conflict, with livestock depredation increasing farther from roads in Hunchun Nature Reserve while peaking at intermediate distances in Kanha National Park (Soh *et al*., 2014; Miller *et al*., 2015). Moreover, human-tiger conflict is often concentrated along forest boundaries and within multiple-use buffer zones where livestock grazing extends into tiger habitat (Kolipaka *et al*., 2017; Dhungana et al., 2017). Although tigers exhibit comparatively low tolerance for sustained, intense anthropogenic disturbance (Harihar *et al*., 2009; Sunarto *et al*., 2012), connected forest landscapes and high wild-prey availability can facilitate movement through human-modified landscapes (Reddy *et al*., 2019; Struebig *et al*., 2018). Human-tiger conflict may therefore peak at intermediate levels of anthropogenic pressure, where disturbance remains sufficiently low for tiger persistence while livestock remain an accessible prey option (Miller *et al*., 2015; Rostro-García *et al*., 2016).

Bioclimatic and hydrological associations with conflict have generally reflected warmer and drier conditions, with higher livestock depredation reported under lower precipitation, higher temperatures, and greater proximity to water sources (Naha, Sathyakumar and Rawat, 2018; Ramesh *et al*., 2020; Sharma *et al*., 2020; Silwal *et al*., 2024). Warmer and drier conditions can alter vegetation productivity and prey distributions while reducing surface-water availability (Abrahms, 2021; Dong and Anderson, 2022). During water scarcity, wildlife, livestock and human activities may therefore concentrate around persistent water sources, increasing spatial overlap and opportunities for conflict (Abade *et al*., 2018; Beattie *et al*., 2020; Senft *et al*., 1987).

Topography further shapes human-carnivore conflict patterns by influencing carnivore occurrence while constraining human activity and livestock grazing patterns (Rather, Kumar and Khan, 2020; Reddy *et al*., 2019). Specifically, gentler slopes have been associated with increased livestock depredation across landscapes (Lamichhane *et al*., 2023). Though relatively understudied, aspect also has distinct implications for conflict patterns, as increased solar exposure on southern slopes has been associated with increases in productivity and foraging by both livestock and wild species, thereby influencing large carnivore distributions (Cheng *et al*., 2024; Lamichhane *et al*., 2023; Letro *et al*., 2022; Soh et al., 2014a; Zeng *et al*., 2010). Elevation, however, has shown variable and often non-linear impact on conflict patterns. For example, human-leopard conflict increased with elevation in the Annapurna Conservation Area, while being concentrated at lower and intermediate elevations in the Kangchenjunga Landscape and Pauri Garhwal, likely reflecting landscape-specific variation in livestock grazing, forest fragmentation, and human land use (Lamichhane *et al*., 2023; Naha, Sathyakumar and Rawat, 2018; Sharma *et al*., 2020). This highlights the need for a comprehensive and integrated analysis of the ecological, environmental, and antropogenic drivers of human-carnivore conflict across heterogeneous landscapes.

However, most studies of human-carnivore conflict patterns remain restricted to individual landscapes or protected areas, single spatial scales, and background sampling methodologies. This limits inference across broader ecological and anthropogenic gradients and multiscale landscape processes, while inadequately accounting for spatial sampling bias (Bommel, Badry and Ford, 2020; Davoli et al., 2022; McGarigal et al., 2016). A regional, multi-synthesis can address this limitation by encompassing broader ecological and anthropogenic gradients, multiple spatial scales, and alternative background-sampling strategies.

## 2. Method

### 2.1 Literature Survey and Conflict Records

A structured literature review identified studies examining spatial patterns and drivers of human-tiger and human-leopard conflict across India, Nepal, Bhutan, and Bangladesh. The literature search covered Google Scholar, Web of Science, and Scopus, using keywords such as “human-tiger conflict,” “human-leopard conflict,” “livestock depredation,” and “spatial patterns of conflict.” A total of 100 unique peer-reviewed publications, preprints, and governmental and NGO technical reports, published until 2025, were screened. For data extraction, conflict records were restricted to direct incidents of conflict, including incidents of livestock depredation, human attacks and retaliatory killings. Precise conflict locations were obtained either from coordinates reported in published datasets or extracted by georeferencing and digitizing published maps in QGIS. Records reported only by locality or as conflict hotspots were excluded. Records were therefore extracted only from 34 publications that provided individual records at appropriate spatial and temporal resolution. Of these, 17 publications provided HLC data, 13 had HTC data, while four covered both. The initial dataset consisted of 4,485 HLC and 3,196 HTC conflict incidents. However, the final dataset was reduced to conflict incidents recorded from 2000 onwards due to limited accurate predictor variable data prior 2000. The final dataset had 4,310 and 2,536 records.

**Figure 1.**
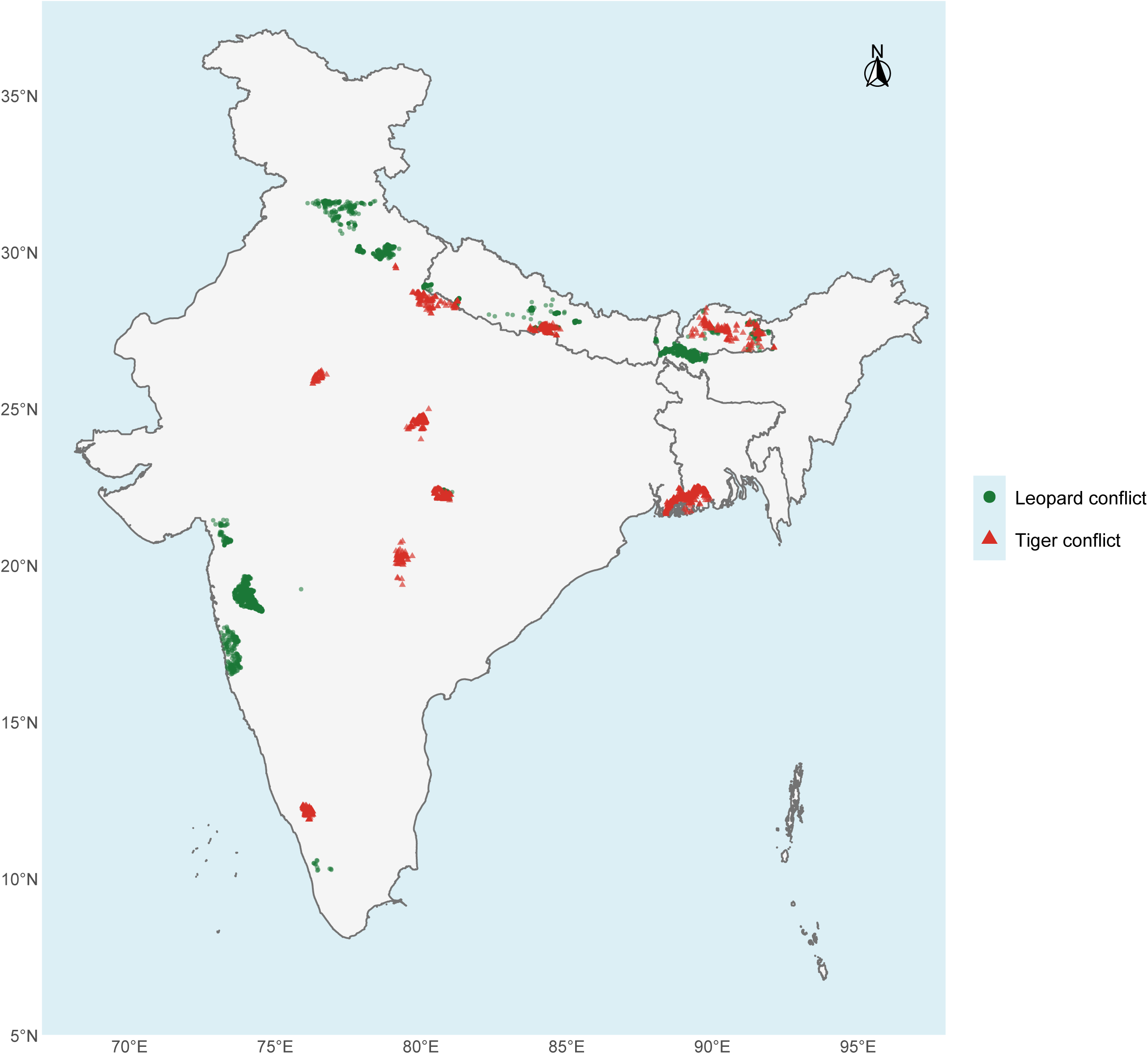
Spatial distribution of human-leopard and human-tiger conflict records across Bangladesh, Bhutan, India, and Nepal from 2000 onwards. The final dataset comprised 4,310 human-leopard conflict (green) and 2,491 human-tiger conflict (red) records. Conflict records included human attacks, livestock depredation and retaliatory killings.

### 2.2 Background Sampling Methodology

Recorded conflict locations were compared with background locations representing potentially available conflict environments without assuming true absence (Barbet-Massin *et al*., 2012; Phillips *et al*., 2009). Three background-sampling strategies - accessible-area, bias-corrected and uniform-random - were evaluated to represent alternative assumptions regarding human accessibility and spatial sampling bias and to assess the sensitivity of model performance to background selection (Fourcade *et al*., 2014; Hanberry, He and Palik, 2012).

Background sampling was restricted to within 50 km of conflict locations to retain relevant landscape variation while excluding distant environments (Hazen *et al*., 2021; VanDerWal *et al*., 2009). Within this extent, a 5-km exclusion zone around conflict incidents was applied to reduce local spatial dependence between conflict and background samples. Background points were generated at 250-m resolution in **Google Earth Engine (Gorelick et al., 2017)**, with each sampling strategy initially targeting 40,000 locations across the full dataset. This target was approximately ten times the size of the larger conflict dataset, standardizing sampling effort while representing available environmental conditions (Barbet-Massin *et al*., 2012). After restricting conflict and background records to 2000 onwards, the HTC background-to-conflict ratio increased to approximately 12:1, whereas the HLC ratio remained relatively unchanged. HTC background records were therefore randomly subsampled without replacement, separately for each sampling method, to achieve a 10:1 ratio. All eligible conflict records were retained, and each eligible background record had an equal probability of selection within its respective method, yielding 25,360 background points per method. A fixed random seed (123) was set before processing the three HTC datasets to ensure reproducibility.

In the accessible area method, background sampling was restricted to areas accessible to human and livestock activity. This reduced the inclusion of areas where few records may reflect limited opportunities for human-carnivore interactions rather than lower conflict risk (Phillips *et al*., 2009). Locations within 5 km of settlements or 2 km of roads were classified as accessible, consistent with conflict patterns reported in central Indian tiger landscapes (Kolipaka *et al*., 2017; Miller, Jhala and Schmitz, 2016). Settlements were identified using GHS-SMOD, with all settlement classes included (Pesaresi and Politis, 2026), while all GRIP4 road classes were retained (Meijer *et al*., 2018). These criteria formed a binary accessibility mask, from which backgrounds were sampled with equal probability.

The bias-corrected method addressed the greater probability that incidents in accessible locations would be encountered and reported (Baker, Maclean and Gaston, 2024; Phillips *et al*., 2009). Accessibility was represented continuously using exponential distance-decay weights for the same GHSL settlements and GRIP4 roads:

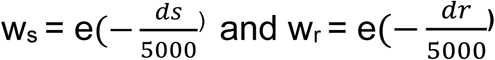

where *w*_s_ and *w*_r_ are the settlement and road weights, and *d*_s_ and *d*_r_ represent the respective Euclidean distances in meters (Baker, Maclean and Gaston, 2024; Fullman *et al*., 2021; Lazenby *et al*., 2021). The 5-km decay scale represented declining accessibility over distances relevant to human and livestock activity (Kolipaka *et al*., 2017; Miller, Jhala and Jena, 2015; Miller, Jhala and Schmitz, 2016). The weights were combined and normalized to a 0-1 bias surface, with a minimum weight of 0.05 permitting selection in less accessible areas. Backgrounds were sampled probabilistically from this surface.

The uniform-random method assigned equal selection probability to all eligible pixels within the study extent. This represented landscape conditions without assumptions regarding accessibility or reporting bias and provided a baseline for evaluating the spatially constrained strategies (Hazen *et al*., 2021).

### 2.3 Predictor Selection and Spatial Scales

Candidate predictors were selected *a priori* from literature on human-tiger and human-leopard conflict and the ecology, habitat use and distribution of both species. Predictors represented landscape structure, anthropogenic disturbance, climate, topography and hydrology. To account for scale-dependent relationships, landscape, topographic and hydrological variables were quantified within 1-, 2-and 5-km circular extents around conflict and background locations (Jackson and Fahrig, 2014; Miller *et al*., 2015). Table A1 summarizes definitions, data sources, and spatial scales for all 33 candidate predictors, and Appendix S1 provides detailed extraction procedures.

To extract the spatial correlates for the predictor variables, remote sensing data were processed using **Google Earth Engine (GEE)** [insert reference/citation]. Spatial data layers including [list variables, e.g., NDVI, elevation, land cover] were accessed and processed using GEE’s cloud-computing platform. The analysis relied entirely on publicly available, open-source secondary datasets…"

Landscape composition and configuration can influence conflict by shaping carnivore distribution, movement, prey availability and exposure to human activity (Fahrig, 2007; McIntyre and Wiens, 1999). Land-cover proportions, Shannon’s Diversity Index, forest patch density, edge density and mean patch area were calculated within the three circular buffers around each conflict and background location using the 30-m Global Land Analysis and Discovery Land Cover and Land Use Change dataset (Potapov *et al*., 2022). Distance to forest was extracted separately at each location. Records were matched to the most recent GLAD product available at or before the observation year from 2000, 2005, 2010, 2015 or 2020. Mean baseline canopy cover and cumulative tree-cover loss were calculated within the three spatial extents using the 30-m Hansen Global Forest Change dataset v1.12 (Hansen *et al*., 2013). Baseline canopy cover represented conditions in 2000, whereas cumulative loss was calculated to the exact observation year. Mean annual NDVI was similarly calculated within each spatial extent from 30-m Landsat Collection 2 Level-2 surface-reflectance imagery matched to the exact observation year (Tucker, 1979; Vermote *et al*., 2016).

Anthropogenic conditions were represented by human population density, built-up cover and distance to settlements as measures of human presence and settlement intensity; road density and distance to roads as measures of infrastructure and accessibility, while livestock density was included as a measure of domestic-prey availability. These factors can alter carnivore occurrence and movement and consequently influence spatial overlap with people and livestock (Naha *et al*., 2020; Rostro-García *et al*., 2016; Sunarto *et al*., 2012). Built-up cover and road density were quantified within the three spatial extents, whereas population and livestock densities and distances to settlements and roads were extracted at each location. Population density was obtained from the approximately 1-km Gridded Population of the World v4 dataset and matched to the 2000, 2005, 2010, 2015 or 2020 product (CIESIN, 2017). Settlement proximity was derived from the 100-m GHS-BUILT-S dataset and matched to the 2000 or 2015 layer (Pesaresi and Politis, 2026). Proximity to roads and road density were derived from the temporally static Global Roads Inventory Project dataset (Meijer *et al*., 2018). Livestock density was obtained from approximately 10-km Gridded Livestock of the World products and matched to the 2010, 2015 or 2020 layer (FAO, 2024, 2026; Gilbert *et al*., 2018). Mean elevation, slope, northness and eastness were calculated within the three spatial extents using the 30-m Copernicus DEM GLO-30 dataset (Amatulli *et al*., 2018; European Space Agency, 2024). Temporal matching was unnecessary because these variables were treated as temporally static.

Hydrological conditions can influence carnivore movements and concentrate wild and domestic prey (Miller, Jhala and Jena, 2015; Ramesh *et al*., 2020; Silwal *et al*., 2024). Therefore, distance to the nearest surface-water pixel was extracted at each location, while mean water occurrence and seasonality were calculated at the three spatial scales using the 30-m Joint Research Center Global Surface Water dataset v1.4. (Pekel *et al*., 2016).

Finally, climatic conditions can influence conflict by altering resource availability, carnivore distributions, and spatial overlap with people and livestock. Six variables associated with carnivore habitat suitability or conflict were extracted at each conflict or background location from the approximately 1-km WorldClim v2.1 dataset: isothermality, maximum temperature of the warmest month, minimum temperature of the coldest month, annual precipitation and precipitation of the wettest and driest quarters (Baral *et al*., 2023; Fick and Hijmans, 2017; Karki *et al*., 2025; Mukul *et al*., 2019).

### 2.4 Statistical Analysis

Human-tiger and human-leopard conflict were modeled separately using binomial generalized linear models with a logit link via the base stats package in R version 4.4.1 (R Core Team, 2024). Conflict and background locations were coded as one and zero, respectively, and models were fitted for each of the three background-sampling methods. Models were evaluated using predictors at 1, 2 and 5km spatial scales. Predictive performance was assessed using five-fold spatial block cross-validation with block sizes of 10, 20, 30, 40 and 50 km^2^, repeated across 100 random fold allocations (Roberts et al., 2017; Valavi et al., 2019) implemented via the blockCV package (Valavi et al., 2019).

Predictor preprocessing and selection were conducted independently within each training fold to prevent information leakage using the caret package (Kuhn, 2008). Missing values were median-imputed. Predictors with zero or near-zero variance or no finite observations were excluded. For predictor pairs with absolute Pearson correlations of at least 0.70, the variable with the greater mean absolute correlation with the remaining predictors was removed to preferentially retain the variable with lower overall collinearity (Dormann et al., 2012; Zuur, Ieno and Elphick, 2010). Predictors were standardized using training-set parameters, and variables were iteratively removed until all variance inflation factors were below five using the car package (Fox and Weisberg, 2019).

Performance was evaluated using PR-AUC, ROC-AUC, Brier score and log-loss. PR-AUC was calculated using a trapezoidal numerical integration of the precision-recall curve mapping across all probability thresholds, while ROC-AUC was computed using the pROC package (Robin et al., 2011). PR-AUC was prioritized because of class imbalance (Gneiting and Raftery, 2007; Saito and Rehmsmeier, 2015). Mean performance across background-sampling methods was used to select the optimal predictor scale and cross-validation block size. The methods were then compared at this spatial configuration. All model configurations are reported in Table S4. Final models were refitted to the full datasets, and effects are reported as odds ratios with 95% confidence intervals. Complete HLC and HTC model outputs are provided in Tables S2 and S3. Permutation importance was calculated as the mean reduction in test-fold ROC-AUC across 30 permutations and cross-validation folds, then averaged across methods in which each predictor was retained.

## 3. Results

### 3.1. Effects of Background Sampling on Predictive Performance

For both species, model performance was evaluated across 15 spatial configurations for each of the three background-sampling methods, comprising all combinations of three landscape spatial scales and five spatial cross-validation block sizes, resulting in 45 evaluated models for both species (Table S4; Fig. S1). The 5-km predictor scale and 10-km cross-validation block size yielded the highest predictive discriminatory ability averaged across the three background-sampling methods for both HLC (PR-AUC = 0.253; ROC-AUC = 0.731) and HTC (PR-AUC = 0.485; ROC-AUC = 0.854). This spatial configuration was retained as the optimal combination for the remaining analysis.

HTC models consistently showed greater predictive discrimination than HLC models across all three background-sampling methods at this optimal configuration (Fig. 2). For HLC, the AA method produced the highest model performance (PR-AUC = 0.288; ROC-AUC = 0.748), followed by UR (PR-AUC = 0.248; ROC-AUC = 0.733) and BC (PR-AUC = 0.222; ROC-AUC = 0.713). Meanwhile, HTC model performance was even more consistent across background sampling methods, with PR-AUC ranging between 0.477 and 0.491 and ROC-AUC ranging from 0.853 to 0.856.

**Figure 2.**
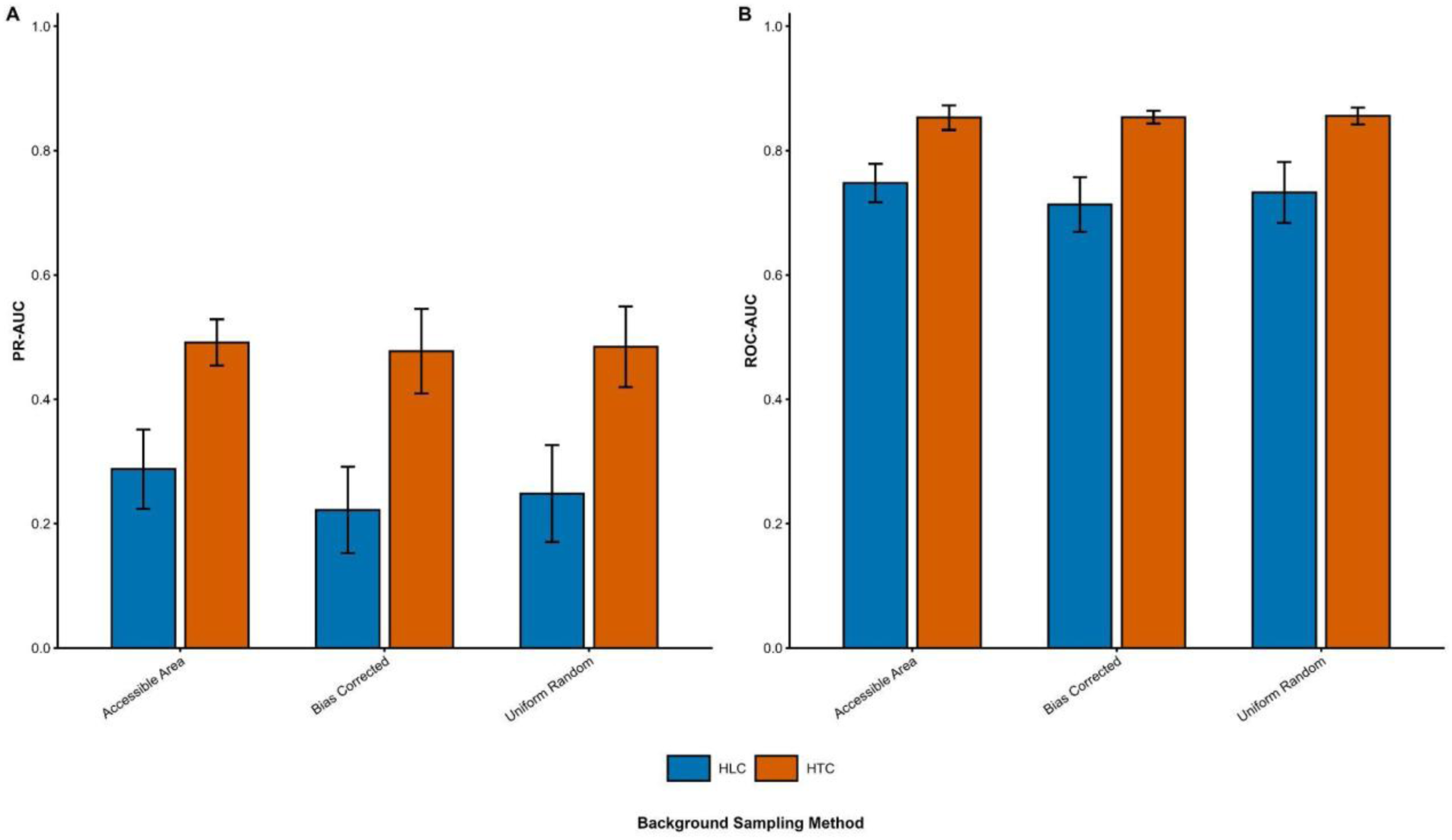
HTC models showed higher predictive performance across all three background sampling methods than HLC, though models of both species were highly consistent. Predictive performance of binomial generalized linear models for (A) human-leopard conflict (HLC) and (B) human-tiger conflict (HTC) was evaluated using precision-recall area under the curve (PR-AUC) and receiver operating characteristic area under the curve (ROC-AUC). Models were evaluated at the optimal spatial configuration of a 5-km predictor extent and 10-km spatial cross-validation block size. Bars show mean performance across five cross-validation folds, and error bars indicate ±1 SD. HLC PR-AUC ranged from 0.222 to 0.288 and ROC-AUC from 0.713 to 0.748 across background-sampling methods. HTC PR-AUC ranged from 0.477 to 0.491 and ROC-AUC from 0.853 to 0.856 across background-sampling methods.

### 3.2 Landscape Predictor Importance and Spatial Correlates of Conflict

#### 3.2.1. Permutation Importance

The relative permutation importance of retained predictors differed considerably between species at the optimal spatial configuration (Fig. 3). Permutation importance was calculated as the mean reduction in test-fold ROC-AUC following 30 permutations of each retained predictor, then averaged across cross-validation folds and the background-sampling methods in which the predictor was retained (Section 2.4).

**Figure 3.**
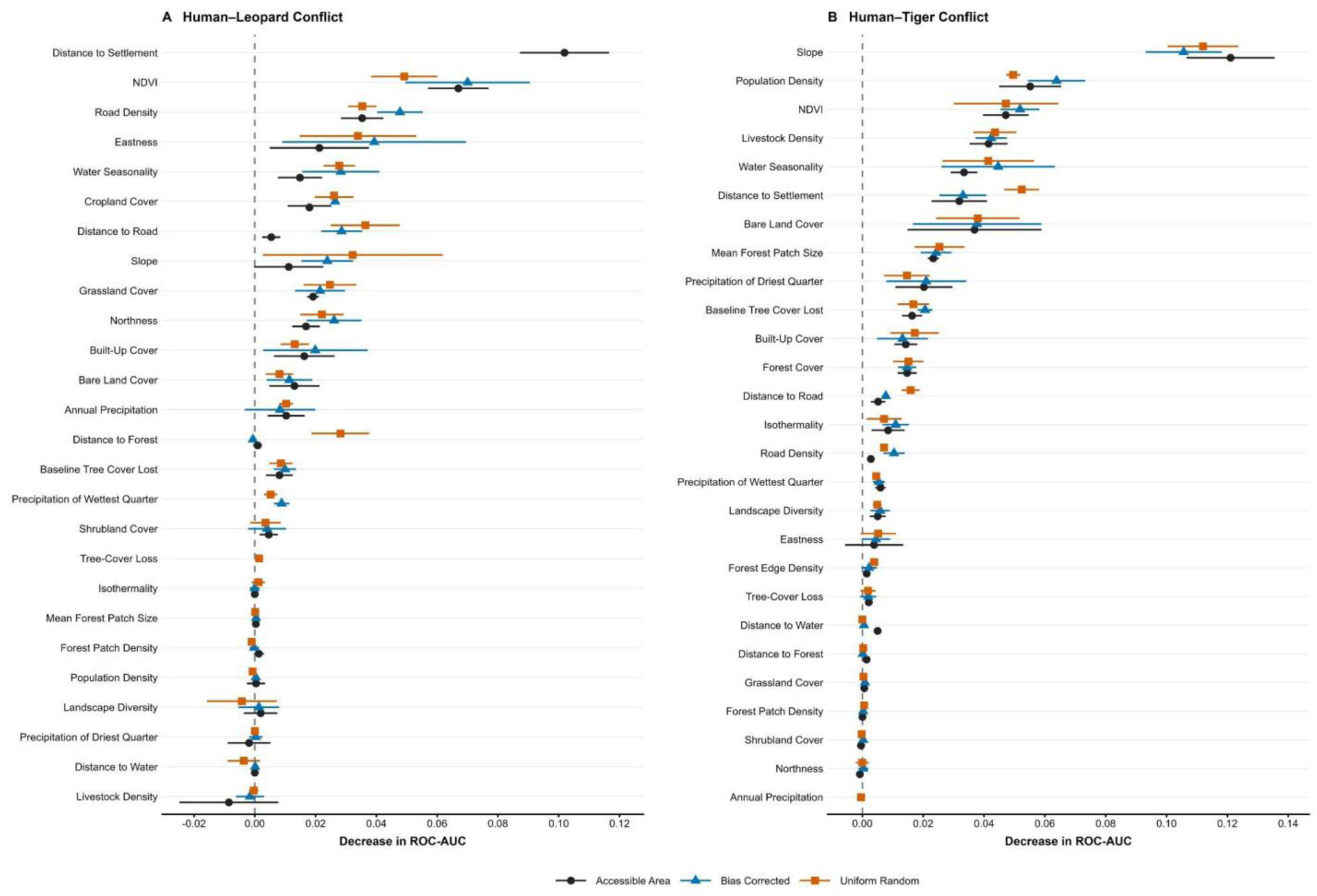
NDVI and road density had the highest mean permutation importance for HLC among variables evaluated under all three background-sampling methods, while slope contributed the most to predictive discrimination for HTC by a substantial margin. Permutation importance was calculated for (A) human-leopard conflict (HLC) and (B) human-tiger conflict (HTC) at the optimal spatial configuration. Permutation importance was quantified as the reduction in test-fold ROC-AUC following predictor permutation (ΔROC-AUC), with larger values indicating greater contributions to predictive discrimination. Points and horizontal error bars represent the mean and ±1 SD of ΔROC-AUC across cross-validation folds within the AA (black), BC (blue), and UR (orange) methods.

Specifically, for HLC, NDVI had the greatest mean permutation importance among predictors retained under all three background-sampling methods (mean ΔROC-AUC = 0.062). This was followed by road density (0.039), eastness (0.031), water seasonality (0.024), cropland cover (0.024), distance to roads (0.023), slope (0.022), grassland cover (0.022), northness (0.022), and built-up cover (0.016). Distance to settlements had the largest mean importance overall (0.102), but was retained only under AA sampling. All remaining HLC predictors had mean ΔROC-AUC values of 0.011 or lower.

Meanwhile, for HTC, slope had substantially greater permutation importance than any other predictor (mean ΔROC-AUC = 0.113), followed by human population density (0.056), NDVI (0.049), livestock density (0.042), water seasonality (0.040), distance to settlements (0.039), bare-ground cover (0.038), mean forest patch area (0.024), and precipitation of the driest quarter (BIO17; 0.019). All remaining HTC predictors had mean ΔROC-AUC values of 0.018 or lower.

#### 3.2.2. Spatial Correlates of Human-Leopard Conflict

The effects of predictor variables on HLC occurrence were largely consistent across background-sampling methods. Of the 33 candidate predictors, 22 were retained across all three methods. Among these, 20 displayed consistent effect directions, of which 15 were statistically significant under every sampling method (Table S2; Fig. S2).

Across all three background-sampling methods, HLC probability was negatively associated with road density (AA: OR = 0.53, 95% CI [0.49, 0.57], *p < 0.05*; BC: OR = 0.49, 95% CI [0.45, 0.53], *p < 0.05*; UR: OR = 0.54, 95% CI [0.50, 0.59], *p < 0.05*; Fig. 4A), but positively associated with built-up cover (AA: OR = 1.32, 95% CI [1.27, 1.38], *p < 0.05*; BC: OR = 1.29, 95% CI [1.24, 1.34], *p < 0.05*; UR: OR = 1.25, 95% CI [1.20, 1.30], *p < 0.05*; Fig. 4B) and Shannon’s Diversity Index (SHDI; AA: OR = 1.23, 95% CI [1.18, 1.29], *p < 0.05*; BC: OR = 1.18, 95% CI [1.13, 1.23], *p < 0.05*; UR: OR = 1.07, 95% CI [1.03, 1.13], *p* = 0.0026; Fig. 4C). HLC probability also decreased significantly with a greater proportion of baseline tree cover lost (AA: OR = 0.74, 95% CI [0.70, 0.79], *p < 0.05*; BC: OR = 0.78, 95% CI [0.74, 0.83], *p < 0.05*; UR: OR = 0.79, 95% CI [0.75, 0.83], *p < 0.05*; Fig. 4D). Conversely, HLC probability was positively associated with indicators of vegetation productivity and precipitation, including NDVI (AA: OR = 2.27, 95% CI [2.09, 2.46], *p < 0.05*; BC: OR = 2.03, 95% CI [1.89, 2.17], *p < 0.05*; UR: OR = 1.86, 95% CI [1.71, 2.02], *p < 0.05*; Fig. 4E) and annual precipitation (AA: OR = 1.23, 95% CI [1.17, 1.30], *p < 0.05*; BC: OR = 1.23, 95% CI [1.16, 1.29], *p < 0.05*; UR: OR = 1.21, 95% CI [1.15, 1.27], *p < 0.05*; Fig. 4F). The reported odds ratios represent the change in odds associated with a one-standard-deviation increase in each continuous predictor.

**Figure 4.**
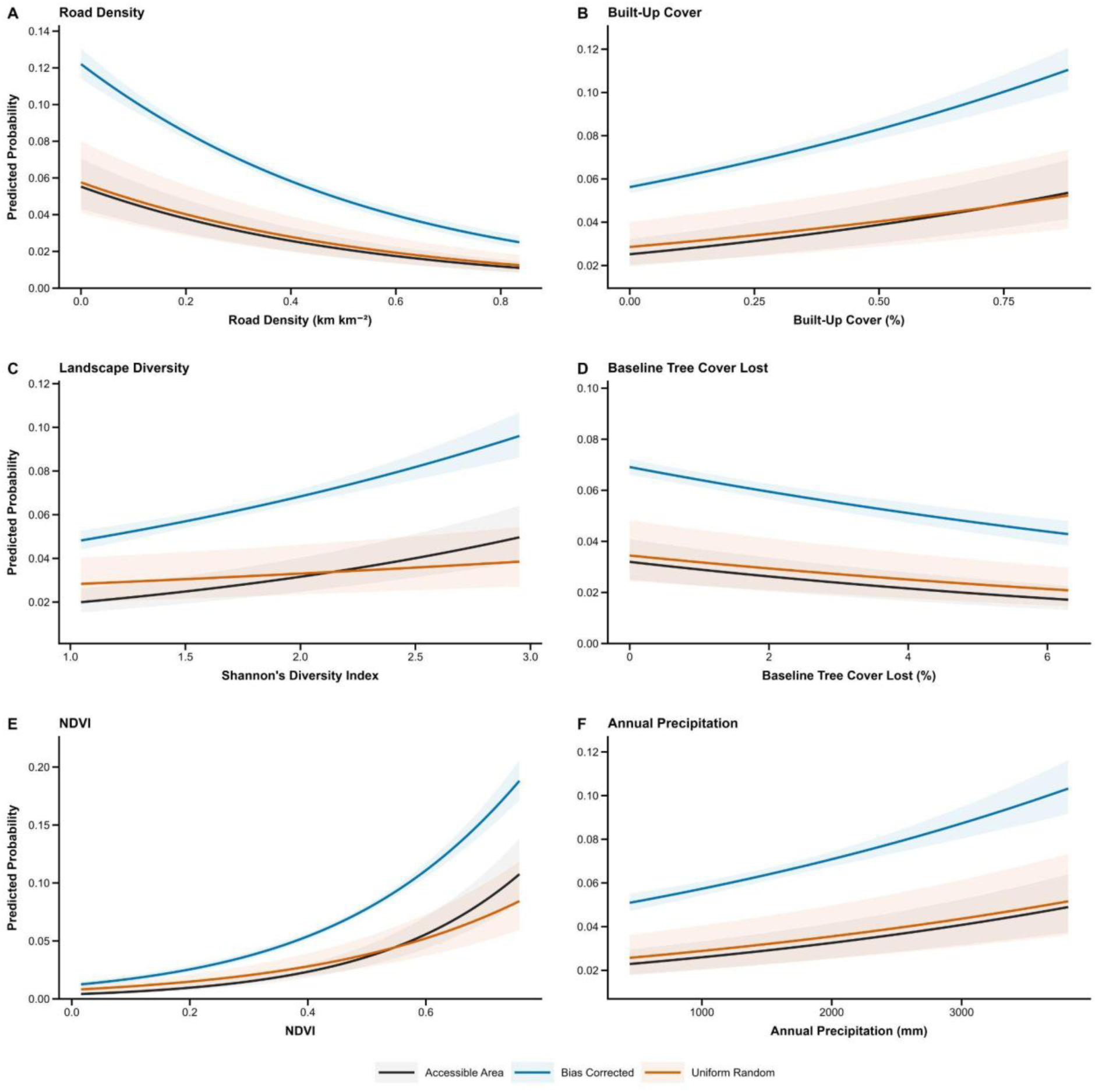
Human-leopard conflict probability increased with built-up cover, landscape heterogeneity, productivity, and annual precipitation, but decreased with road density and tree-cover loss. The partial-response curves show variation in predicted HLC probability with (A) road density, (B) built-up cover, (C) Shannon’s Diversity Index, (D) proportion of baseline tree cover lost, (E) NDVI, and (F) annual precipitation across the three background-sampling methods. Each variable was derived from full binomial GLMs evaluated under the optimal spatial configuration, with all other continuous predictors held at their standardized means. Lines represent predicted probabilities, and shaded bands indicate 95% confidence intervals.

Similarly, across all three background-sampling methods, HLC probability was significantly higher with greater forest patch density, shrubland cover, and eastness, and significantly lower with greater bare-ground and grassland cover, northness, distance to roads, water seasonality, and slope (Table S2; Fig. S2). Distance to forest, livestock density, mean forest patch size, precipitation of the driest quarter, and human population density showed consistent effect directions across sampling methods but were not statistically significant under every method. The remaining predictors varied in effect direction, statistical significance, or retention across methods (Table S2; Fig. S2).

#### 3.2.3. Spatial Correlates of Human-Tiger Conflict

The spatial correlates of HTC were also robust to background selection, as 23 of the 33 predictor variables had consistent effect directions across all three methods, including 18 that were consistently significant (Table S3; Fig. S2).

HTC occurrence was negatively associated with livestock density (AA: OR = 0.41, 95% CI [0.36, 0.47], *p < 0.05*; BC: OR = 0.40, 95% CI [0.35, 0.46], *p < 0.05*; UR: OR = 0.40, 95% CI [0.35, 0.45], *p < 0.05*; Fig. 5A) and human population density (AA: OR = 0.09, 95% CI [0.07, 0.12], *p < 0.05*; BC: OR = 0.03, 95% CI [0.02, 0.04], *p < 0.05*; UR: OR = 0.08, 95% CI [0.06, 0.10], *p < 0.05*; Fig. 5B). Precipitation during the driest quarter was positively associated with HTC occurrence (AA: OR = 1.73, 95% CI [1.63, 1.84], *p < 0.05*; BC: OR = 1.72, 95% CI [1.62, 1.83], *p < 0.05*; UR: OR = 1.58, 95% CI [1.48, 1.68], *p < 0.05*; Fig. 5C). HTC occurrence declined with increasing mean forest patch size (AA: OR = 0.24, 95% CI [0.15, 0.41], *p < 0.05*; BC: OR = 0.26, 95% CI [0.16, 0.41], *p < 0.05*; UR: OR = 0.20, 95% CI [0.11, 0.35], *p < 0.05*; Fig. 5D) and increased with forest cover (AA: OR = 1.71, 95% CI [1.60, 1.82], *p < 0.05*; BC: OR = 1.72, 95% CI [1.61, 1.84], *p < 0.05*; UR: OR = 1.83, 95% CI [1.70, 1.97], *p < 0.05*; Fig. 5E). Conversely, SHDI was negatively associated with HTC occurrence (AA: OR = 0.76, 95% CI [0.72, 0.80], *p < 0.05*; BC: OR = 0.74, 95% CI [0.70, 0.78], *p < 0.05*; UR: OR = 0.76, 95% CI [0.72, 0.81], *p < 0.05*; Fig. 5F).

**Figure 5.**
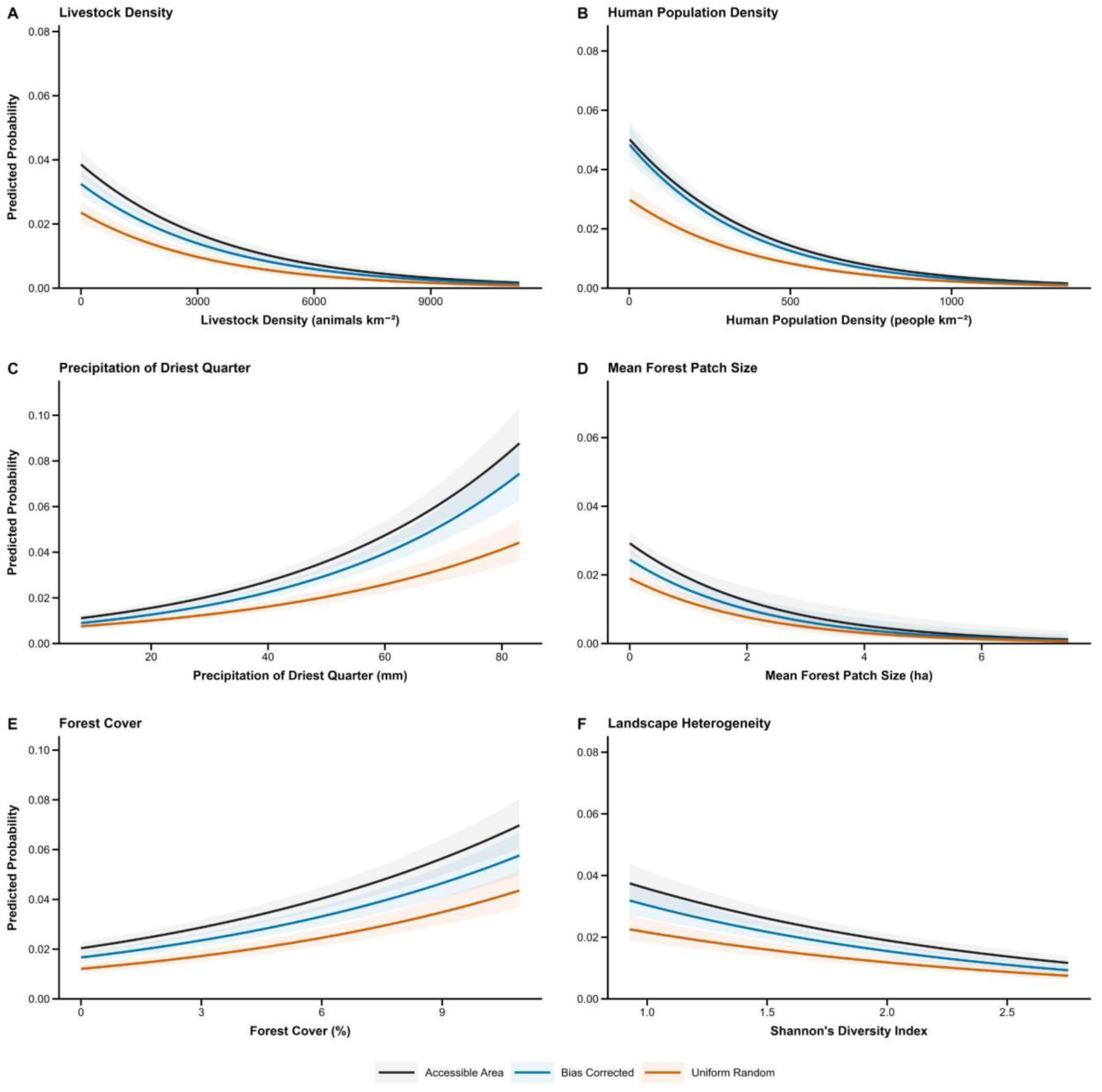
Predicted human-tiger conflict probability increased with precipitation during the driest quarter and forest cover, but declined with livestock density, human population density, mean forest patch size and landscape heterogeneity. Partial-response curves are shown for (A) livestock density, (B) human population density, (C) precipitation during the driest quarter, (D) mean forest patch size, (E) forest cover and (F) Shannon’s Diversity Index under the three background-sampling methods. Predictions were derived from full binomial GLMs at the optimal spatial configuration. Each predictor was varied across the central 95% of its observed range while other continuous predictors were held at their standardized means. Lines show predicted probabilities, and shaded bands indicate 95% confidence intervals.

HTC showed consistent, significant positive associations with bare-ground cover, grassland cover, NDVI, eastness, northness and water seasonality. Meanwhile, significant negative associations were observed with built-up cover, isothermality, distance to settlements, road density and slope. Other predictors varied in direction, significance or retention across background-sampling methods (Table S3).

## 4. Discussion

This study provides a comprehensive assessment of the spatial drivers of human-wildlife conflict across South Asia, highlighting distinct species-specific conflict patterns. Specifically, HLC was concentrated in productive, heterogeneous, human-modified landscapes, while HTC was associated with more fragmented landscapes with lower anthropogenic pressure. Furthermore, bioclimatic conditions and surface water availability had distinct, and often contrasting, influences on conflict patterns for each species. Conversely, topographic conditions had a uniform influence on conflict risk patterns across both species, highlighting shared implications of topography for species occupancy, habitat use, and anthropogenic activity.

The spatial correlates of human-wildlife conflict were highly robust for both species, with model predictive discrimination remaining broadly consistent across alternative background-sampling methods. This suggests that model performance was robust to alternative sampling assumptions within the ecologically and anthropogenically heterogeneous landscapes represented in the study. Despite strong sampling bias concentrating conflict incident data within only a subset of the regional distribution of both species, the consistently high model performance across background sampling methods highlights the transferability across the wider species ranges (Kramer-Schadt et al., 2013; Phillips et al., 2009).

Furthermore, for both species, model predictive discrimination was highest at the 5-km spatial scale. Although finer spatial scales capture ecological conditions at individual conflict sites, the lower predictive performance at these scales may reflect substantial variation in local conditions across the ecologically heterogeneous landscapes represented in this study. Higher model performance at broader spatial scales may therefore reflect more accurate characterization of landscape correlates of carnivore occurrence and movement, and thus the ecological conditions associated with human-carnivore conflict (Fahrig, 2007; McGarigal et al., 2016). Previous studies have found antropogenic disturbance and habitat structure to influence tigers and leopards at broader spatial scales (Hussain *et al*., 2026; Rather, Kumar and Khan, 2020).

Evaluating different spatial cross-validation block sizes across background sampling methods and predictor spatial scales for both species showed model performance peaking at the 10-km block size across species, then declining with broader block sizes. This implies that analysis adequately accounted for spatial autocorrelation and the underlying ecological determinants of conflict at finer scales (Roberts et al., 2017; Valavi et al., 2019). Lastly, HTC models consistently outperformed HLC models across all combinations of background sampling methods, spatial scales, and cross-validation block sizes. This differential performance reflects the contrasting ecological niche breadths of both species, further reinforced by landscape structure’s differing influence on conflict patterns (Abrego and Ovaskainen, 2023).

HLC was associated with productive, heterogeneous human-modified landscapes, while HTC was concentrated in fragmented forest landscapes with moderate anthropogenic disturbance. Specifically, conflict was positively correlated with forest patch density, NDVI, and shrubland cover but negatively associated with grassland and bare-ground cover across background methods. These patterns align with previous studies that found HLC incidents to increase in landscapes with productive and structurally complex vegetation (Lamichhane *et al*., 2023; Naha *et al*., 2020; Ramesh *et al*., 2020; Yadav, Behera and Chatakonda, 2024). In human-modified landscapes, productive and structurally complex vegetation may support leopard persistence by providing concealment for movement and ambush hunting (Kshettry, Vaidyanathan and Athreya, 2017; Naha *et al*., 2020, 2021). While HTC was also associated with increased forest cover and productivity (Chatterjee *et al*., 2022; Soh et al., 2014a), this potentially reflects the narrower ecological niche of tigers, which are significantly more dependent on intact forest ecosystems (Miller; Sunarto *et al*., 2012; Hussain *et al*., 2026).

Surprisingly, tree cover loss was negatively correlated with HLC probability. Increased habitat loss and degradation were expected to increase livestock depredation due to the reduction in wild-prey densities (Inskip and Zimmermann, 2009; Bhattacharya *et al*., 2025). However, the reduced incidence of conflict in areas with higher tree loss may reflect the concentration of conflict in more productive and structurally complex vegetation. However, this highlights the importance of integrating both wild prey densities and more temporally dynamic measures of land-use change for a more mechanistic understanding of conflict patterns (Inskip and Zimmermann, 2009. This is particularly essential as this study found landscape fragmentation and anthropogenic disturbance to have contrasting impacts on conflict patterns.

Across background sampling methods, HLC increased with landscape heterogeneity, built-up cover, and proximity to roads. In heterogeneous landscapes, the ecological plasticity of leopards allows for the increased exploitation of anthropogenic resources, particularly closer to settlements and human-dominated land uses (Naha *et al*., 2020; Yadav, Behera and Chatakonda, 2024). Both increased built-up area and proximity to roads, have been associated with increased overlap of leopards, livestock, and humans, increasing risk of negative interactions (Athreya *et al*., 2014; Odden *et al*., 2014; Sharma *et al*., 2020; Silwal *et al*., 2024). However, road density was consistently negatively correlated with HLC. This suggests that while landscape heterogeneity and anthropogenic land use may increase accessibility to livestock, leopards may still show some sensitivity to higher levels of anthropogenic disturbance and fragmentation (Abade et al., 2018; Hinde et al., 2023).

Contrastingly, this study found HTC to reduce in more heterogeneous landscapes and with increased anthropogenic pressure, such as increased human population, built-up cover and road densities. This may reflect tigers’ lower tolerance for intensive habitat modification and anthropogenic disturbance (Sunarto *et al*., 2012; Hussain *et al*., 2026).

However, HTC increased closer to settlements and with higher forest-edge density and smaller mean forest patches. These patterns suggest conflict may therefore be concentrated in landscapes where forest fragmentation expands spatial interface between critical tiger habitat (Zanuari et al., 2024). In landscapes with higher tiger densities, fragmented forests may facilitate the natural dispersal of tigers through human-modified landscape, potentially increasing the opportunities for negative interactions with humans and livestock (Chatterjee et al., 2022; Zanuari et al., 2024).. Moreover, this study found HTC to increase with greater grassland cover. Both wild herbivores and livestock herders utilize these ecosystems as grazing pastures, particularly near settlements. near forests, thereby increasing opportunities for depredation (Kolipaka *et al*., 2017; Sharma, Hope and Neupane, 2024). These patterns suggest that HTC is concentrated in landscapes with sufficient forest habitat to support tiger persistence, with substantial spatial overlap with areas utilized by people and livestock.

Surprisingly, livestock density was strongly negatively correlated with HTC, contrary to the expectation that increased livestock availability would increase conflict risk (Khanal *et al*., 2020; Poudel *et al*., 2026). However, coarse livestock density data may not accurately reflect livestock exposure and availability near tiger habitats and conflict hotspots. Furthermore, livestock depredation intensity may be influenced more by grazing practices, guarding, enclosure quality, and wild-prey availability than by livestock abundance alone (Kolipaka *et al*., 2017; Malviya and Krishnamurthy, 2022; Sijapati *et al*., 2021).

Bioclimatic and hydrological conditions had contrasting implications for HTC and HLC patterns. HTC increased with precipitation during the driest quarter, water seasonality, and proximity to water sources. Dry-season rainfall may help maintain landscape productivity and resource availability across seasons, while persistent water sources may concentrate wildlife, livestock, and people around shared resources, thereby increasing conflict opportunities (Pekel *et al*., 2016; Ramesh *et al*., 2020; Chatterjee *et al*., 2022; Miller, Jhala and Schmitz, 2016). In contrast, HLC was negatively correlated with water seasonality, suggesting that seasonal resource constraints likely intensified conflict by concentrating wildlife, livestock, and humans (Abrahms, 2021).

Lastly, topography more consistently influenced conflict patterns for both species. For example, conflict probability was lower on more rugged slopes for both species as these landscapes have been associated with reduced accessibility and limited human and livestock movement (Lamichhane *et al*., 2023a; Ramesh *et al*., 2020). In contrast, this study could not identify consistent correlations between elevation and conflict, potentially reflecting the multidimensional interactions of elevation and other spatial determinants of conflict (Lamichhane *et al*., 2023; Naha, Sathyakumar and Rawat, 2018; Sharma *et al*., 2020). Aspect showed more contrasting patterns between species. Specifically, HLC was concentrated on southerly aspects, a pattern consistent with human-leopard interactions in western Nepal, while HTC was concentrated on northern slopes (Lamichhane *et al*., 2023). Although eastness was positively associated with conflict for both species, further research is needed to clarify the ecological mechanisms underlying these topographic patterns (Letro *et al*., 2021).

While this study identifies distinct spatial correlates of conflict for both species, a more mechanistic understanding of the intricate landscape processes driving conflict is limited by several factors. For example, the coarse temporal resolution of historical conflict records limited the dynamic land-use change and degradation metrics. Moreover, future analyses should also integrate standardized government compensation records, carnivore occupancy or density, wild-prey abundance, and socioeconomic demographics of victims.

## Supporting information

Supplementary Materials and Results

## 6. Declarations

### CRediT Author Statement

Anish Banerjee: Conceptualization, Methodology, Software, Validation, Formal analysis, Investigation, Data curation, Writing - original draft, Writing - review & editing, Visualization.

## Generative AI Disclosure

During the preparation of this work, the author used ChatGPT (OpenAI; GPT-6 Astra) to help develop and debug JavaScript code to extract predictor variable data from Google Earth Engine and R code for statistical analyses. The author reviewed, tested, and edited the outputs and takes full responsibility for the manuscript content.

## Data Availability

The supplementary materials list the publications used to compile conflict records (Table S1), describe the predictor datasets and extraction procedures (Appendix S1 and Table A1), and report model coefficients and predictive performance (Figures S1-S2 and Tables S2-S4). The Google Earth Engine JavaScript and R scripts used for background sampling, predictor extraction, and statistical analyses are available at [GitHub repository URL]. The compiled conflict dataset will be made available upon reasonable request.

