## Supplementary material for "Understanding Spatial Patterns of Human-Tiger and Human-Leopard Conflict across South Asia using a Multiscale Regional Approach": S2

### Table S2. Complete HLC model coefficients and consistency across background-sampling methods.

Human–leopard conflict. Full binomial GLMs; three background-sampling methods.

| Predictor | Method | Coefficient [95% CI] | SE | Odds ratio [95% CI] | p-value | Methods retaining predictor | Consistent direction across all three | Significant across all three |
|---|---|---|---|---|---|---|---|---|
| Annual Precipitation | AA | 0.209 [0.157, 0.262] | 0.027 | 1.233 [1.170, 1.299] | <0.001 | 3 | Yes: positive | Yes |
| Annual Precipitation | BC | 0.203 [0.153, 0.254] | 0.026 | 1.225 [1.165, 1.289] | <0.001 | 3 | Yes: positive | Yes |
| Annual Precipitation | UR | 0.191 [0.141, 0.242] | 0.026 | 1.211 [1.151, 1.274] | <0.001 | 3 | Yes: positive | Yes |
| Bare Land Cover | AA | -0.347 [-0.403, -0.290] | 0.029 | 0.707 [0.668, 0.748] | <0.001 | 3 | Yes: negative | Yes |
| Bare Land Cover | BC | -0.275 [-0.328, -0.222] | 0.027 | 0.760 [0.721, 0.801] | <0.001 | 3 | Yes: negative | Yes |
| Bare Land Cover | UR | -0.216 [-0.270, -0.163] | 0.027 | 0.806 [0.764, 0.850] | <0.001 | 3 | Yes: negative | Yes |
| Baseline Tree Cover Lost | AA | -0.296 [-0.357, -0.235] | 0.031 | 0.744 [0.700, 0.790] | <0.001 | 3 | Yes: negative | Yes |
| Baseline Tree Cover Lost | BC | -0.246 [-0.305, -0.187] | 0.030 | 0.782 [0.737, 0.829] | <0.001 | 3 | Yes: negative | Yes |
| Baseline Tree Cover Lost | UR | -0.237 [-0.292, -0.181] | 0.028 | 0.789 [0.747, 0.834] | <0.001 | 3 | Yes: negative | Yes |
| Built-Up Cover | AA | 0.277 [0.236, 0.318] | 0.021 | 1.319 [1.266, 1.375] | <0.001 | 3 | Yes: positive | Yes |
| Built-Up Cover | BC | 0.252 [0.212, 0.291] | 0.020 | 1.286 [1.237, 1.338] | <0.001 | 3 | Yes: positive | Yes |
| Built-Up Cover | UR | 0.221 [0.181, 0.260] | 0.020 | 1.247 [1.199, 1.297] | <0.001 | 3 | Yes: positive | Yes |
| Cropland Cover | AA | -1.584 [-2.962, -0.205] | 0.703 | 0.205 [0.052, 0.814] | 0.024 | 2 | Not assessable | Not assessable |
| Cropland Cover | BC | Not retained | — | — | — | 2 | Not assessable | Not assessable |
| Cropland Cover | UR | -3.575 [-5.504, -1.645] | 0.984 | 0.028 [0.004, 0.193] | <0.001 | 2 | Not assessable | Not assessable |
| Distance to Forest | AA | -0.291 [-0.403, -0.179] | 0.057 | 0.748 [0.668, 0.837] | <0.001 | 3 | Yes: negative | No |

### Table S2. Complete HLC model coefficients and consistency across background-sampling methods.

Human–leopard conflict. Full binomial GLMs; three background-sampling methods.

| Predictor | Method | Coefficient [95% CI] | SE | Odds ratio [95% CI] | p-value | Methods retaining predictor | Consistent direction across all three | Significant across all three |
|---|---|---|---|---|---|---|---|---|
| Distance to Forest | BC | -0.007 [-0.046, 0.032] | 0.020 | 0.993 [0.955, 1.032] | 0.717 | 3 | Yes: negative | No |
| Distance to Forest | UR | -1.145 [-1.288, -1.003] | 0.073 | 0.318 [0.276, 0.367] | <0.001 | 3 | Yes: negative | No |
| Distance to Road | AA | -0.205 [-0.268, -0.142] | 0.032 | 0.815 [0.765, 0.868] | <0.001 | 3 | Yes: negative | Yes |
| Distance to Road | BC | -0.612 [-0.705, -0.519] | 0.048 | 0.542 [0.494, 0.595] | <0.001 | 3 | Yes: negative | Yes |
| Distance to Road | UR | -0.754 [-0.864, -0.644] | 0.056 | 0.470 [0.421, 0.525] | <0.001 | 3 | Yes: negative | Yes |
| Distance to Settlement | AA | -2.239 [-2.420, -2.058] | 0.092 | 0.107 [0.089, 0.128] | <0.001 | 1 | Not assessable | Not assessable |
| Distance to Settlement | BC | Not retained | — | — | — | 1 | Not assessable | Not assessable |
| Distance to Settlement | UR | Not retained | — | — | — | 1 | Not assessable | Not assessable |
| Distance to Water | AA | -0.008 [-0.046, 0.029] | 0.019 | 0.992 [0.955, 1.029] | 0.656 | 3 | No | No |
| Distance to Water | BC | 0.019 [-0.015, 0.053] | 0.017 | 1.019 [0.986, 1.054] | 0.264 | 3 | No | No |
| Distance to Water | UR | 0.012 [-0.030, 0.053] | 0.021 | 1.012 [0.970, 1.055] | 0.585 | 3 | No | No |
| Eastness | AA | 0.368 [0.317, 0.420] | 0.026 | 1.445 [1.372, 1.521] | <0.001 | 3 | Yes: positive | Yes |
| Eastness | BC | 0.443 [0.394, 0.492] | 0.025 | 1.557 [1.482, 1.635] | <0.001 | 3 | Yes: positive | Yes |
| Eastness | UR | 0.418 [0.368, 0.468] | 0.026 | 1.519 [1.445, 1.597] | <0.001 | 3 | Yes: positive | Yes |
| Forest Patch Density | AA | 0.261 [0.185, 0.337] | 0.039 | 1.298 [1.203, 1.401] | <0.001 | 3 | Yes: positive | Yes |
| Forest Patch Density | BC | 0.073 [0.034, 0.112] | 0.020 | 1.076 [1.034, 1.119] | <0.001 | 3 | Yes: positive | Yes |

### Table S2. Complete HLC model coefficients and consistency across background-sampling methods.

Human–leopard conflict. Full binomial GLMs; three background-sampling methods.

| Predictor | Method | Coefficient [95% CI] | SE | Odds ratio [95% CI] | p-value | Methods retaining predictor | Consistent direction across all three | Significant across all three |
|---|---|---|---|---|---|---|---|---|
| Forest Patch Density | UR | 0.093 [0.043, 0.143] | 0.025 | 1.098 [1.044, 1.154] | <0.001 | 3 | Yes: positive | Yes |
| Grassland Cover | AA | -0.410 [-0.487, -0.333] | 0.039 | 0.664 [0.614, 0.717] | <0.001 | 3 | Yes: negative | Yes |
| Grassland Cover | BC | -0.385 [-0.460, -0.309] | 0.038 | 0.681 [0.631, 0.734] | <0.001 | 3 | Yes: negative | Yes |
| Grassland Cover | UR | -0.470 [-0.547, -0.392] | 0.040 | 0.625 [0.579, 0.676] | <0.001 | 3 | Yes: negative | Yes |
| Isothermality | AA | -0.058 [-0.112, -0.004] | 0.028 | 0.943 [0.894, 0.996] | 0.034 | 3 | No | No |
| Isothermality | BC | -0.040 [-0.091, 0.010] | 0.026 | 0.960 [0.913, 1.010] | 0.117 | 3 | No | No |
| Isothermality | UR | 0.052 [-0.000, 0.105] | 0.027 | 1.054 [1.000, 1.110] | 0.050 | 3 | No | No |
| Landscape Diversity | AA | 0.209 [0.163, 0.256] | 0.024 | 1.233 [1.177, 1.292] | <0.001 | 3 | Yes: positive | Yes |
| Landscape Diversity | BC | 0.163 [0.121, 0.205] | 0.021 | 1.177 [1.129, 1.228] | <0.001 | 3 | Yes: positive | Yes |
| Landscape Diversity | UR | 0.071 [0.025, 0.118] | 0.024 | 1.074 [1.025, 1.125] | 0.003 | 3 | Yes: positive | Yes |
| Livestock Density | AA | 0.008 [-0.027, 0.042] | 0.018 | 1.008 [0.974, 1.043] | 0.660 | 3 | Yes: positive | No |
| Livestock Density | BC | 0.070 [0.036, 0.103] | 0.017 | 1.072 [1.036, 1.109] | <0.001 | 3 | Yes: positive | No |
| Livestock Density | UR | 0.058 [0.025, 0.091] | 0.017 | 1.060 [1.025, 1.096] | <0.001 | 3 | Yes: positive | No |
| Mean Forest Patch Size | AA | 0.062 [0.023, 0.102] | 0.020 | 1.064 [1.024, 1.107] | 0.002 | 3 | Yes: positive | No |
| Mean Forest Patch Size | BC | 0.014 [-0.025, 0.053] | 0.020 | 1.014 [0.975, 1.054] | 0.478 | 3 | Yes: positive | No |
| Mean Forest Patch Size | UR | 0.011 [-0.030, 0.051] | 0.021 | 1.011 [0.971, 1.052] | 0.607 | 3 | Yes: positive | No |

**Table S2. Complete HLC model coefficients and consistency across background-sampling methods.**

Human–leopard conflict. Full binomial GLMs; three background-sampling methods.

| Predictor | Method | Coefficient [95% CI] | SE | Odds ratio [95% CI] | p-value | Methods retaining predictor | Consistent direction across all three | Significant across all three |
|---|---|---|---|---|---|---|---|---|
| NDVI | AA | 0.819 [0.738, 0.900] | 0.041 | 2.268 [2.091, 2.459] | <0.001 | 3 | Yes: positive | Yes |
| NDVI | BC | 0.706 [0.636, 0.775] | 0.035 | 2.025 [1.889, 2.171] | <0.001 | 3 | Yes: positive | Yes |
| NDVI | UR | 0.621 [0.539, 0.702] | 0.042 | 1.860 [1.715, 2.018] | <0.001 | 3 | Yes: positive | Yes |
| Northness | AA | -0.294 [-0.334, -0.254] | 0.020 | 0.746 [0.716, 0.776] | <0.001 | 3 | Yes: negative | Yes |
| Northness | BC | -0.309 [-0.348, -0.269] | 0.020 | 0.734 [0.706, 0.764] | <0.001 | 3 | Yes: negative | Yes |
| Northness | UR | -0.313 [-0.353, -0.272] | 0.021 | 0.732 [0.702, 0.762] | <0.001 | 3 | Yes: negative | Yes |
| Population Density | AA | -0.083 [-0.162, -0.004] | 0.040 | 0.920 [0.850, 0.996] | 0.040 | 3 | Yes: negative | No |
| Population Density | BC | -0.078 [-0.162, 0.006] | 0.043 | 0.925 [0.850, 1.006] | 0.068 | 3 | Yes: negative | No |
| Population Density | UR | -0.038 [-0.096, 0.020] | 0.030 | 0.963 [0.908, 1.021] | 0.202 | 3 | Yes: negative | No |
| Precipitation of Driest Quarter | AA | -0.098 [-0.155, -0.042] | 0.029 | 0.906 [0.856, 0.959] | <0.001 | 3 | Yes: negative | No |
| Precipitation of Driest Quarter | BC | -0.043 [-0.097, 0.011] | 0.028 | 0.958 [0.908, 1.011] | 0.122 | 3 | Yes: negative | No |
| Precipitation of Driest Quarter | UR | -0.014 [-0.068, 0.041] | 0.028 | 0.987 [0.934, 1.041] | 0.624 | 3 | Yes: negative | No |
| Road Density | AA | -0.640 [-0.716, -0.564] | 0.039 | 0.527 [0.489, 0.569] | <0.001 | 3 | Yes: negative | Yes |
| Road Density | BC | -0.717 [-0.798, -0.635] | 0.042 | 0.488 [0.450, 0.530] | <0.001 | 3 | Yes: negative | Yes |
| Road Density | UR | -0.610 [-0.684, -0.536] | 0.038 | 0.543 [0.505, 0.585] | <0.001 | 3 | Yes: negative | Yes |
| Shrubland Cover | AA | 0.208 [0.155, 0.261] | 0.027 | 1.231 [1.168, 1.298] | <0.001 | 3 | Yes: positive | Yes |

### Table S2. Complete HLC model coefficients and consistency across background-sampling methods.

Human–leopard conflict. Full binomial GLMs; three background-sampling methods.

| Predictor | Method | Coefficient [95% CI] | SE | Odds ratio [95% CI] | p-value | Methods retaining predictor | Consistent direction across all three | Significant across all three |
|---|---|---|---|---|---|---|---|---|
| Shrubland Cover | BC | 0.183 [0.129, 0.237] | 0.027 | 1.201 [1.138, 1.267] | <0.001 | 3 | Yes: positive | Yes |
| Shrubland Cover | UR | 0.154 [0.099, 0.209] | 0.028 | 1.166 [1.104, 1.232] | <0.001 | 3 | Yes: positive | Yes |
| Slope | AA | -0.312 [-0.377, -0.246] | 0.034 | 0.732 [0.686, 0.782] | <0.001 | 3 | Yes: negative | Yes |
| Slope | BC | -0.360 [-0.419, -0.301] | 0.030 | 0.698 [0.658, 0.740] | <0.001 | 3 | Yes: negative | Yes |
| Slope | UR | -0.422 [-0.485, -0.359] | 0.032 | 0.656 [0.616, 0.699] | <0.001 | 3 | Yes: negative | Yes |
| Water Seasonality | AA | -0.300 [-0.346, -0.254] | 0.024 | 0.741 [0.707, 0.776] | <0.001 | 3 | Yes: negative | Yes |
| Water Seasonality | BC | -0.369 [-0.415, -0.324] | 0.023 | 0.691 [0.660, 0.723] | <0.001 | 3 | Yes: negative | Yes |
| Water Seasonality | UR | -0.385 [-0.432, -0.338] | 0.024 | 0.681 [0.649, 0.713] | <0.001 | 3 | Yes: negative | Yes |

Notes: AA, Accessible Area; BC, Bias Corrected; UR, Uniform Random; SE, standard error; CI, confidence interval.

Coefficients describe the change in log odds of recorded conflict relative to background locations per one-standard-deviation increase in a continuous predictor, conditional on the other predictors.

Confidence intervals are 95% Wald intervals. Odds ratios and their intervals were obtained by exponentiating coefficients and confidence limits. For the intercept, exponentiation gives baseline odds.

Reported p-values were retained; missing p-values were calculated using a two-sided Wald z test where possible. Values below 0.001 are displayed as <0.001.

Statistical significance was assessed at $p < 0.05$ without adjustment for multiple testing. Consistency and significance were assessed from unrounded values.

Direction is consistent when all three methods have estimable coefficients with the same nonzero sign. Significance across all three methods is assessed separately from directional consistency.

Not retained indicates that the predictor is absent from the saved model. A dash indicates that no model estimate applies; NA denotes an unavailable quantity for a retained term.

### Table S2. Complete HLC model coefficients and consistency across background-sampling methods.

Human–leopard conflict. Full binomial GLMs; three background-sampling methods.

| Predictor | Method | Coefficient [95% CI] | SE | Odds ratio [95% CI] | p-value | Methods retaining predictor | Consistent direction across all three | Significant across all three |
|---|---|---|---|---|---|---|---|---|

Not assessable indicates insufficient information across all three methods. Intercept consistency is not assessed. Candidate inventories are included when supplied; otherwise, rows cover predictors retained in at least one method.
