## Supplementary material for "Understanding Spatial Patterns of Human-Tiger and Human-Leopard Conflict across South Asia using a Multiscale Regional Approach": S1

| Reference ID | Species | Country | Study Area | Keywords Used |
|---|---|---|---|---|
| 1 | Tiger | India | Sundarbans | Spatial Drivers of Human-Tiger Conflict |
| 2 | Tiger | India | Panna | Livestock Depradation Patterns |
| 3 | Both | India | Multiple | Livestock Depradation Patterns |
| 4 | Tiger | India | Yavatamal | Human-Tiger Conflict |
| 5 | Tiger | India | Corbett | Human-Willdife Conflict Patterns |
| 6 | Tiger | India | Corbett | Spatial Drivers of Human-Tiger Conflict |
| 7 | Tiger | India | Ranthambore | Spatial Drivers of Human-Tiger Conflict |
| 8 | Tiger | India | Dudhwa | Spatial Drivers of Human-Tiger Conflict |
| 9 | Both | India | Kanha | Livestock Depradation Patterns |
| 10 | Tiger | India | Nagarhole | Spatial Drivers of Human-Tiger Conflict |
| 11 | Tiger | Nepal | Chitwan | Spatial Drivers of Human-Tiger Conflict |
| 12 | Both | Bhutan | National | Livestock Depradation Patterns |
| 13 | Tiger | India | Panna | Human-Tiger Conflict |
| 14 | Tiger | India | Panna | Human-Tiger Conflict |
| 15 | Tiger | Nepal | Chitwan | Human-Tiger Conflict |
| 16 | Tiger | Bangladesh | Sundarbans | Spatial Drivers of Human-Tiger Conflict |
| 17 | Tiger | Bangladesh | Sundarbans | Human-Tiger Conflict |
| 18 | Tiger | Bangladesh | Sundarbans | Livestock Depradation Patterns |
| 19 | Tiger | India | Jim Corbett | Spatial Drivers of Human-Tiger Conflict |
| 20 | Tiger | India | Jim Corbett | Spatial Drivers of Human-Tiger Conflict |
| 21 | Both | Nepal | Khata | Spatial Drivers of Human-Tiger Conflict |
| 22 | Tiger | Nepal | Chitwan | Human-Tiger Conflict |
| 23 | Both | Nepal | Chitwan | Spatial Drivers of Human-Tiger Conflict |
| 24 | Tiger | Nepal | Chitwan | Human-Tiger Conflict |
| 25 | Tiger | Nepal | Chitwan | Human-Tiger Conflict |
| 26 | Tiger | India | Tadoba | Livestock Depradation Patterns |
| 27 | Tiger | India | Nagarhole | Human-Tiger Conflict |
| 28 | Tiger | India | Kanha | Spatial Drivers of Human-Tiger Conflict |
| 29 | Tiger | Nepal | Badria | Human-Tiger Conflict |
| 30 | Tiger | Nepal | Bardia | Human-Tiger Conflict |
| 31 | Tiger | Nepal | Chitwan | Human-Willdife Conflict Patterns |
| 32 | Tiger | India | Ranthambore | Human-Tiger Conflict |

| | | | | |
|---|---|---|---|---|
| 33 | Tiger | Nepal | Khatia | Human-Tiger Conflict |
| 34 | Tiger | Nepal | Bardia | Human-Tiger Conflict |
| 35 | Tiger | Nepal | Chitwan | Human-Tiger Conflict |
| 36 | Tiger | India | Sundarbans | Human-Tiger Conflict |
| 37 | Leopard | India | Ratnagiri | Livestock Depradation Patterns |
| 38 | Leopard | India | Karnataka | Human-Leopard Conflict |
| 39 | Leopard | India | Gujarata | Livestock Depradation Patterns |
| 40 | Leopard | India | Rajasthan | Spatial Drivers of Human-Leopard Conflict |
| 41 | Leopard | India | Karnataka | Human-Leopard Conflict |
| 42 | Leopard | India | Bengal/NE | Human-Leopard Conflict |
| 43 | Leopard | India | Junnar | Spatial Drivers of Human-Leopard Conflict |
| 44 | Leopard | India | Junnar | Human-Leopard Conflict |
| 45 | Leopard | India | Rajaji | Livestock Depradation Patterns |
| 46 | Leopard | India | Kerala-Tamil Nadu | Human-Leopard Conflict |
| 47 | Leopard | Nepal | Shuklaphanta | Spatial Drivers of Human-Leopard Conflict |
| 48 | Leopard | India | Uttharakhand | Human-Leopard Conflict |
| 49 | Leopard | India | Eastern Himalayas | Spatial Drivers of Human-Leopard Conflict |
| 50 | Leopard | Nepal | Chitwan | Spatial Drivers of Human-Leopard Conflict |
| 51 | Leopard | Nepal | Mid-Hills | Spatial Drivers of Human-Leopard Conflict |
| 52 | Leopard | Nepal | Bardia | Spatial Drivers of Human-Leopard Conflict |
| 53 | Leopard | India | Surat | Human-Leopard Conflict |
| 54 | Leopard | India | Jammu and Kashmir | Livestock Depradation Patterns |
| 55 | Leopard | | | Livestock Depradation Patterns |
| 56 | Leopard | India | Bardia | Spatial Drivers of Human-Leopard Conflict |
| 57 | Leopard | Nepal | Aadhikhola | Spatial Drivers of Human-Leopard Conflict |
| 58 | Leopard | India | Kailash | Human-Willdife Conflict Patterns |
| 59 | Leopard | India | GHNP | Livestock Depradation Patterns |
| 60 | Leopard | Nepal | National | Human-Willdife Conflict Patterns |
| 61 | Leopard | Nepal | Baitadi | Human-Leopard Conflict |
| 62 | Leopard | Nepal | National | Spatial Drivers of Human-Leopard Conflict |
| 63 | Leopard | Nepal | Kathmandu | Human-Leopard Conflict |
| 64 | Leopard | India | Maharasthra | Human-Leopard Conflict |

| | | | | |
|---|---|---|---|---|
| 65 | Leopard | India | Pauri Garwhal | Human-Leopard Conflict |
| 66 | Leopard | Nepal | Kathmandu | Human-Leopard Conflict |
| 67 | Leopard | Nepal | Entire | Human-Willdife Conflict Patterns |
| 68 | Leopard | Multiple | Multiple | Livestock Depradation Patterns |
| 69 | Leopard | India | Uttharakhand | Human-Leopard Conflict |
| 70 | Leopard | Nepal | Gandaki Province | Human-Willdife Conflict Patterns |
| 71 | Leopard | Nepal | Pokhara Valley | Livestock Depradation Patterns |
| 72 | Leopard | Nepal | Mid Hill | Human-Willdife Conflict Patterns |
| 73 | Leopard | Nepal | Chitwan | Human-Willdife Conflict Patterns |
| 74 | Leopard | India | Maharasthra | Spatial Drivers of Human-Leopard Conflict |
| 75 | Leopard | India | Maharashtra | Livestock Depradation Patterns |
| 76 | Leopard | Nepal | Shivapuri | Human-Leopard Conflict |
| 77 | Leopard | Nepal | Kathmandu | Livestock Depradation Patterns |
| 78 | Leopard | India | Binsar | Livestock Depradation Patterns |
| 79 | Leopard | Nepal | Gandaki Province | Human-Leopard Conflict |
| 80 | Leopard | India | Western Ghats | Human-Willdife Conflict Patterns |
| 81 | Leopard | Nepal | Gandaki Province | Human-Leopard Conflict |
| 82 | Leopard | Nepal | Annupurma | Livestock Depradation Patterns |
| 83 | Leopard | Nepal | Lalitpur | Human-Leopard Conflict |
| 84 | Leopard | India | Terai | Human-Willdife Conflict Patterns |
| 85 | Leopard | India | Gujarat | Livestock Depradation Patterns |
| 86 | Leopard | India | Kali Tiger Reserve | Human-Willdife Conflict Patterns |
| 87 | Leopard | Nepal | Gorkha | Livestock Depradation Patterns |
| 88 | Leopard | Nepal | Gaurishankar | Human-Leopard Conflict |
| 89 | Leopard | Nepal | Bardia—Banke Complex | Human-Willdife Conflict Patterns |
| 90 | Leopard | India | East and Western Ghats | Livestock Depradation Patterns |
| 91 | Leopard | India | Corbett | Human-Willdife Conflict Patterns |

| | | | | |
|---|---|---|---|---|
| 92 | Leopard | India | Aravalli | Human-Leopard Conflict |
| 93 | Leopard | India/Nepal | Kanchenjunga | Human-Willdife Conflict Patterns |
| 94 | Leopard | India | Valparai | Livestock Depradation Patterns |
| 95 | Leopard | Nepal | Bardia | Human-Willdife Conflict Patterns |
| 96 | Leopard | India | Pir Panjal | Livestock Depradation Patterns |
| 97 | Leopard | Nepal | Chitwan | Livestock Depradation Patterns |
| 98 | Leopard | Bhutan | Jigme Singye Wangchuck National Park | Livestock Depradation Patterns |
| 99 | Leopard | India | Gir | Human-Leopard Conflict |
| 100 | Both | India | Kanha National Park | Human-Willdife Conflict Patterns |
| | | | | 17 |
| | | | | 13 |
| | | | | 4 |

| Publication Year | Type of Source | Data Used | Used? | Type of Data | Data Collection Method |
|---|---|---|---|---|---|
| 2022 | Database | Yes | NA | Raw Data | Raw Data |
| 2022 | Publication | Yes | NA | Raw Data | Raw Data |
| 2018 | Publication | Yes | NA | Raw Data | Raw Data |
| 2020 | Publication | No | Coarse | Vilage Name | NA |
| 2024 | Publication | No | Coarse | Vilage Name | NA |
| 2015 | Publication | No | Coarse | Vilage Name | NA |
| 2014 | Publication | Yes | NA | Published Map | QGIS |
| 2022 | Publication | Yes | NA | Published Map | QGIS |
| 2016 | Publication | Yes | NA | Published Map | QGIS |
| 2025 | Publication | Yes | NA | Published Map | QGIS |
| 2017 | Publication | Yes | NA | Published Map | QGIS |
| 2016 | Publication | Yes | NA | Published Map | QGIS |
| 2024 | Publication | Yes | NA | Published Map | QGIS |
| 2017 | Publication | Yes | NA | Published Map | QGIS |
| 2017 | Publication | Yes | NA | Published Map | QGIS |
| 2015 | Publication | Yes | NA | Published Map | QGIS |
| 2013 | Publication | Yes | NA | Raw Data | Raw Data |
| 2009 | Publication | Yes | NA | Raw Data | Raw Data |
| 2026 | Publication | No | Broad Temporal Range | Published Map | QGIS |
| 2015 | Publication | Yes | NA | Published Map | QGIS |
| 2024 | Publication | Yes | NA | Published Map | QGIS |
| 2008 | Publication | No | Overlap | NA | NA |
| 2020 | Publication | No | Overlap | NA | NA |
| 2025 | Publication | No | Overlap | NA | NA |
| 2016 | Publication | No | Overlap | NA | NA |
| 2013 | Publication | No | Overlap | NA | NA |
| 2025 | Publication | No | Overlap | NA | NA |
| 2015 | Publication | No | Overlap | NA | NA |
| 2014 | Publication | Not Available | NA | NA | NA |
| 2019 | Publication | Not Available | NA | NA | NA |
| 2019 | Publication | Not Available | NA | NA | NA |
| 2025 | Publication | Not Available | NA | NA | NA |

| | | | | | |
|---|---|---|---|---|---|
| 2024 | Publication | Not Available | NA | NA | NA |
| 2025 | Publication | Not Available | NA | NA | NA |
| 2020 | Publication | Not Available | NA | NA | NA |
| 2025 | Publication | Not Available | NA | NA | NA |
| 2011 | Publication | Yes | NA | Published Map | QGIS |
| 2014 | Publication | Not used | | Raw Data | N/A |
| 2021 | Publication | Yes | NA | Published Map | QGIS |
| 2021 | Publication | Not used | Coarse | Raw Data | Raw Data |
| 2023 | Publication | Not used | Coarse | Raw Data | Raw Data |
| 2017 | Publication | Not used | Coarse | Raw Data | Raw Data |
| 2023 | Publication | Yes | NA | Published Map | QGIS |
| 2010 | Publication | Yes | NA | Published Map | QGIS |
| 2023 | Publication | Yes | NA | Published Map | QGIS |
| 2021 | Publication | Yes | NA | Published Map | QGIS |
| 2023 | Publication | Yes | NA | Published Map | QGIS |
| 2020 | Publication | Yes | NA | Raw Data | Raw |
| 2020 | Publication | Yes | NA | Published Map | QGIS |
| 2018 | Publication | Yes | NA | Raw | Raw |
| 2024 | Publication | Yes | NA | Raw | Raw |
| 2023 | Publication | Not used | Broad Temporal Resoultion | Published Map | QGIS |
| 2018 | Publication | Yes | NA | Published Map | QGIS |
| 2023 | Publication | Not used | Congregrate Data | Published Map | QGIS |
| 2018 | Publication | Not used | Pakistan data | Published Map | QGIS |
| 2025 | Publication | Yes | NA | Published Map | QGIS |
| 2021 | Publication | Yes | NA | Published Map | QGIS |
| 2024 | Publication | Not used | Broad Temporal Resolution | Published Map | QGIS |
| 2024 | Publication | Not used | Could not access Open Source | NA | NA |
| 2017 | Publication | Yes | NA | Published Map | QGIS |
| 2014 | Publication | Not used | Coarse Data | Raw | Raw |
| 2023 | Publication | Not used | Hotspot Analysus | Published Map | QGIS |
| 2017 | Publication | Yes | NA | Published Map | QGIS |
| 2020 | Publication | Yes | NA | Published Map | QGIS |

| | | | | | |
|---|---|---|---|---|---|
| 2025 | Publication | Not used | Broad Temporal Range | Published Map | QGIS |
| 2025 | Publication | Not used | Broad Temporal Range | Published Map | QGIS |
| 2016 | Publication | Not Available | NA | NA | NA |
| 2015 | Publication | Not Available | NA | NA | NA |
| 2023 | Publication | Not Available | NA | NA | NA |
| 2022 | Publication | Not Available | NA | NA | NA |
| 2019 | Publication | Not Available | NA | NA | NA |
| 2018 | Publication | Not Available | NA | NA | NA |
| 2024 | Publication | Not Available | NA | NA | NA |
| 2021 | Publication | Not Available | NA | NA | NA |
| 2014 | Publication | Not Available | NA | NA | NA |
| 2019 | Publication | Not Available | NA | NA | NA |
| 2022 | Publication | Not Available | NA | NA | NA |
| 2013 | Publication | Not Available | NA | NA | NA |
| 2023 | Publication | Not Available | NA | NA | NA |
| 2013 | Publication | Not Available | NA | NA | NA |
| 2022 | Publication | Not Available | NA | NA | NA |
| 2023 | Publication | Not Available | NA | NA | NA |
| 2022 | Publication | Not Available | NA | NA | NA |
| 2015 | Publication | Not Available | NA | NA | NA |
| 2023 | Publication | Not Available | NA | NA | NA |
| 2023 | Publication | Not Available | NA | NA | NA |
| 2023 | Publication | Not Available | NA | NA | NA |
| 2024 | Publication | Not Available | NA | NA | NA |
| 2024 | Publication | Not Available | NA | NA | NA |
| 2020 | Publication | Not Available | NA | NA | NA |
| 2025 | Publication | Not Available | NA | NA | NA |

| | | | | | |
|---|---|---|---|---|---|
| 2018 | Publication | Not Available | NA | NA | NA |
| 2020 | Publication | Not Available | NA | NA | NA |
| 2017 | Publication | Not Available | NA | NA | NA |
| 2021 | Publication | Not Available | NA | NA | NA |
| 2022 | Publication | Not Available | NA | NA | NA |
| 2024 | Publication | Not Available | NA | NA | NA |
| 2006 | Publication | Not Available | NA | NA | NA |
| 2022 | Publication | Not Available | NA | NA | NA |
| 2018 | Publication | Not Available | NA | NA | NA |

| Related | Notes |
| --- | --- |
| NA | NA |
| NA | NA |
| https://www.ca | https://www.res |
| NA | NA |
| NA | NA |
| NA | NA |
| NA | NA |
| NA | NA |
| https://onlinelib | NA |
| https://journala | NA |
| NA | NA |
| NA | NA |
| NA | NA |
| NA | NA |
| NA | NA |
| NA | NA |
| NA | NA |
| NA | NA |
| NA | NA |
| NA | NA |
| NA | NA |
| NA | NA |
| NA | NA |
| NA | NA |
| NA | NA |
| NA | NA |
| NA | NA |
| NA | NA |
| NA | NA |
| NA | NA |
| NA | NA |
| NA | NA |
| NA | NA |

| | |
|---|---|
| NA | NA |
| NA | NA |
| NA | NA |
| NA | NA |
| NA | NA |
| NA | NA |
| NA | NA |
| NA | NA |
| NA | NA |
| NA | NA |
| NA | NA |
| NA | NA |
| NA | NA |
| NA | NA |
| NA | NA |
| NA | NA |
| NA | NA |
| NA | NA |
| N/A | N/A |
| NA | NA |
| N/A | N/A |
| N/A | N/A |
| NA | NA |
| NA | NA |
| NA | NA |
| NA | NA |
| NA | NA |
| NA | NA |
| NA | NA |
| NA | NA |
| NA | NA |

| | |
|---|---|
| NA | NA |
| NA | NA |
| NA | NA |
| NA | NA |
| NA | NA |
| NA | NA |
| NA | NA |
| NA | NA |
| NA | NA |
| NA | NA |
| NA | NA |
| NA | NA |
| NA | NA |
| NA | NA |
| NA | NA |
| NA | NA |
| NA | NA |
| NA | NA |
| NA | NA |
| NA | NA |
| NA | NA |
| NA | NA |
| NA | NA |
| NA | NA |
| NA | NA |
| NA | NA |
| NA | NA |
| NA | NA |

| | |
|---|---|
| NA | NA |
| NA | NA |
| NA | NA |
| NA | NA |
| NA | NA |
| NA | NA |
| NA | NA |
| NA | NA |
| NA | NA |

| Citation |
| --- |
| Indian sundarban. Harvard Dataverse. doi:10.7910/dvn/ys0wte. |
| reintroduced tiger (panthera tigris) to predict conflict risk probability. Global Ecology and Conservation, 40, |
| Kushnir, H., Linnell, J.D.C. and Fountain-Jones, N.M. (2019). Species-specific spatiotemporal patterns of leopard, lion and tiger attacks on humans. Journal of Applied Ecology, 56(3), pp.585–593. doi:10.1111/1365- |
| and challenges of changing behaviour of wild animals in human dominated landscape. International Journal of Scientific Research in Biological Sciences, 7(4), pp.71–77. |
| Corbett National Park Ramnagar Uttarakhand, India (from november 2023 to january 2024). International Journal of Science and Research Archive, 11(1), pp.2214–2218. doi:10.30574/ijsra.2024.11.1.0301. |
| scenario. Asian Journal of Science and Technology, 6(10), pp.1925–1927. |
| Characterizing human–tiger conflict in and around Ranthambhore Tiger Reserve, Western India. European Journal of Wildlife Research, 61(2), pp.255–261. doi:10.1007/s10344-014-0895-z. |
| tiger (panthera tigris) interactions in mosaic landscapes around Dudhwa and pilibhit tiger reserves in India. Frontiers in Conservation Science, 3. doi:10.3389/fcosc.2022.999195. |
| for livestock: Implications for mitigating human-carnivore conflict. PLOS ONE, 11(9), p.e0162685. |
| Tiger Reserve, southern part of India: A five-year case study. Journal of Environmental Biology, [online] 46(6), |
| Tigers panthera tigris: Patterns, correlates, and contexts of human–tiger conflict in Chitwan National Park, Nepal. Oryx, 52(1), pp.55–65. doi:10.1017/s0030605316001587. |
| dependence of felid predation risk: Identifying predictors of livestock kills by tiger and leopard in Bhutan. Landscape Ecology, 31(6), pp.1277–1298. doi:10.1007/s10980-015-0335-9. |
| carnivore respond to anthropogenic pressures at fine-scales? Implications for interface area management. PeerJ, [online] 12, pp.e17693-e17693. doi:10.7717/peerj.17693. |
| domestic prey in the diet of reintroduced tigers (panthera tigris) in the livestock-dominated multiple-use forests of Panna Tiger Reserve, India. PLOS ONE, 12(4), p.e0174844. doi:10.1371/journal.pone.0174844. |
| S., Mishra, R., Dhakal, M., Smith, J.L.D. and de Iongh, H.H. (2017). Are conflict-causing tigers different? Another perspective for understanding human-tiger conflict in Chitwan National Park, Nepal. Global Ecology |
| Bangladesh understanding patterns and processes. 27th International Congress for Conservation Biology. . |
| D. (2013). Human–tiger conflict in context: Risks to lives and livelihoods in the Bangladesh Sundarbans. Human Ecology: An interdiscplinary Journal, 41(2), pp.169–186. doi:10.1007/s10745-012-9556-6. |
| tdge of the Bangladesh Sundarbans - Technical Report. Wildlife Trust of Bangladesh. |
| Rawat, M.S., Rawat, A. and Rawat, P.K. (2026). Adverse ecological changes along transitional buffer zone of Jim Corbett National Park increasing human-tiger conflict risk: Geospatial approach. Environmental Monitoring and Assessment, 198(1). doi:10.1007/s10661-025-14936-0. |
| –human conflict in Corbett Tiger Reserve (CTR,) India: Based on the genetic analysis. Wildlife Biology in Practice, 11(1). doi:10.2461/wbp.2015.11.1. |
| wildlife interactions: Evidence from forest edges of Khata Corridor, Bardia, western Nepal. Nepalese Journal of Zoology, 8(2), pp.40–47. doi:10.3126/njz.v8i2.74930. |
| killing tigers in Chitwan National Park, Nepal. Biological Conservation, 141(12), pp.3069–3078. |
| Ruda, A., Kolejka, J. and Silwal, T. (2020). Spatial concentrations of wildlife attacks on humans in Chitwan National Park, Nepal. Animals, 10(1), p.153. doi:10.3390/ani10010153. |
| Tiger (Panthera Tigris) Conflict: Status and Trend in the Chitwan National Park, Nepal. International Journal of Research and Scientific Innovation, XII(V), pp.500–511. doi:10.51244/ijrsi.2025.12050044. |
| of compensation payments and tiger removals in Chitwan National Park, Nepal. Tropical Conservation Science, 9(2), pp.776–787. doi:10.1177/194008291600900213. |
| Large carnivore attacks on humans in central India: A case study from the tadoba-andhari tiger reserve. Oryx, [online] 47(2), pp.221–227. doi:10.1017/S0030605311001803. |
| resource management: A human-tiger conflict case study from Nagarahole Tiger Reserve. Asian Journal of Education and Social Studies, 51(9), pp.649–662. doi:10.9734/ajess/2025/v51i92404. |
| tigers: Implications of spatial grain for modeling predation risk to mitigate human-carnivore conflict. Ecology and Evolution, 5(6), pp.1354–1367. doi:10.1002/ece3.1440. |
| Bhattarai, B.R. and Fischer, K. (2014). Human–tiger panthera tigris conflict and its perception in Bardia National Park, Nepal. Oryx, 48(4), pp.522–528. doi:10.1017/s0030605313000483. |
| Lessons from Bardia and Chitwan National Parks, Nepal. European Journal of Wildlife Research, 65(3). doi:10.1007/s10344-019-1270-x. |
| Mishra, R. and de Iongh, H.H. (2019). Contribution of buffer zone programs to reduce human-wildlife impacts: The case of the Chitwan National Park, Nepal. Human Ecology, [online] 47(1), pp.95–110. doi:10.1007/s10745- |
| wildlife conflict: Assessing the impact of land use land cover change in Ranthambhore Tiger Reserve, India. Environmental Research Communications. doi:10.1088/2515-7620/ade229. |

| |
|---|
| coexistence in the forests of the Khata Corridor, Bardiya, Nepal. Oryx, [online] pp.1–9.<br>doi:10.1017/s0030605323001849. |
| Human–tiger conflict and community responses in Bardiya National Park, Nepal. Frontiers in Conservation<br>Science, [online] 6. doi:10.3389/fcosc.2025.1623056. |
| mitigation efforts in Madi Valley of Chitwan District, Nepal. Journal of Forest and Natural Resource<br>Management, 2(1), pp.37–49. doi:10.3126/jfnrm.v2i1.40219. |
| Pathak, S., Mukherjee, J., Sen, A. and Choudry, A. (2025). Whose habitat? Exploring human-tiger conflict in<br>the riskscapes of the Indian sundarbans. Journal of Political Ecology, 32(1). doi:10.2458/jpe.5111. |
| response attitudes of people affected with livestock depredation by leopards in Ratnagiri district, Maharashtra<br>(India) . Journal of Bombay Natural Hisotry Society, 103(1), pp.18–23. |
| Using media reports to examine leopard distribution, depredation, and management practices outside protected<br>areas in southern India. PLOS ONE, 10(11), p.e0142647. doi:10.1371/journal.pone.0142647. |
| in Vansda Taluka, South Gujarat. Journal of Entomology and Zoology Studies, 9(4), pp.218–226. |
| interactions based on media reports in northwestern India. Journal of Threatened Taxa, 13(6),<br>pp.18453–18478. doi:10.11609/jott.7244.13.6.18453-18478. |
| Karnataka: A review from media reports. Journal of Global Ecology and Environment, pp.16–29. |
| human-leopard interactions in north-eastern India. PLOS ONE, 12(5), p.e0177013. |
| analysis in Junnar Forest Division of Pune Forest Circle, Maharashtra, India. European Journal of Wildlife<br>Research, 70(1). doi:10.1007/s10344-023-01755-2. |
| conflict with leopards in human-dominated landscapes of India. Conservation Biology, [online] 25(1), |
| the forest-human interphase area of northern India. 1. doi:https://doi.org/10.22541/au.168320350.05133789/v1. |
| Govind, S.K. and Jayson, E.A. (2021). Human-leopard conflict on the Kerala-Tamil Nadu border, Southern<br>India. Journal of the Bombay Natural History Society (JBNHS), 118(0). doi:10.17087/jbnhs/2021/v118/148822. |
| and effectiveness of mitigation in Shuklaphanta National Park, Nepal. PLoS ONE, 18(4), p.e0282654.<br>doi:10.1371/journal.pone.0282654. |
| deterrents to reduce livestock losses to leopard attacks within a multiple-use landscape of the himalayan |
| (2020). Landscape predictors of human–leopard conflicts within multi-use areas of the himalayan region.<br>Scientific Reports, 10(1). doi:10.1038/s41598-020-67980-w. |
| B.P. and de Iongh, H.H. (2018). Spatio-temporal patterns of attacks on human and economic losses from<br>wildlife in Chitwan National Park, Nepal. PLOS ONE, 13(4), p.e0195373. doi:10.1371/journal.pone.0195373. |
| (2024). Identifying risk zones and landscape features that affect common leopard depredation. PeerJ, 12,<br>p.e17497. doi:10.7717/peerj.17497. |
| Sandhya Dhakal, Rimal, S., Prashant Paudel and Shrestha, A. (2023). Spatio-temporal patterns of livestock<br>predation by leopards in Bardia National Park, Nepal. Land, 12(6), pp.1156–1156. doi:10.3390/land12061156. |
| Dominated Landscape and Mitigation Measures: A Case Study from Surat District, Gujarat - Technical Report. |
| western himalayan region of Jammu and Kashmir: Temporal adherence in predator’s choice. Tropical Ecology,<br>65(1), pp.26–31. doi:10.1007/s42965-022-00290-6. |
| Loss of biological diversity leads to trouble. Biodiversity and Conservation, 27(11), pp.2795–2813. |
| leopard (panthera pardus) attacks on livestock in buffer zone of Bardia National Park, Nepal. Asian Journal of<br>Research in Zoology, [online] 8(1), pp.89–107. doi:10.9734/ajriz/2025/v8i1186. |
| attacks: A case study from Aadhikhola Rural Municipality, Syangja. Forestry: Journal of Institute of Forestry,<br>Nepal, [online] 18(01), pp.81–89. doi:10.3126/forestry.v18i01.41759. |
| Hussain, A., Sarkar, M.S., Adhikari, B.S., Sathyakumar, S. and Rawat, G.S. (2024). The sacred and the wild:<br>Unpacking human-wildlife interaction in Indian part of Kailash Sacred Landscape. Journal of Wildlife Science,<br>1(3), pp.111–121. doi:10.63033/jwls.tgwi5580. |
| Bhardwaj, S., Thakur, S., Singh, A.P. and De, K. (2024). Livestock depredation pattern by common leopard: A<br>case study from great himalayan national park conservation area. National Academy Science Letters, 48(3),<br>pp.373–377. doi:10.1007/s40009-024-01470-9. |
| configuration work as indicators of human–wildlife conflict? Evidences from human death and injury by wildlife<br>attacks in Nepal. Ecological Indicators, 80, pp.74–83. doi:10.1016/j.ecolind.2017.04.037. |
| Special Issues on Biodiversity Conservation Efforts in Nepal. Kathmandu, Nepal: Department of National Parks |
| and anthropogenic drivers of leopard (panthera pardus fusca) attack occurrence on humans in Nepal. People<br>and nature, 5(6), pp.1977–1988. doi:10.1002/pan3.10536. |
| Thesis] Available at:<br>https://elibrary.tucl.edu.np/JQ99OgQIizUxyjI9nB0on9OyLkqsGIf4/api/core/bitstreams/7f99cfcf-abac-4677-835c- |
| The impact of leopards ( panthera pardus ) on livestock losses and human injuries in a human-use landscape<br>in Maharashtra, India. PeerJ, 8, p.e8405. doi:10.7717/peerj.8405. |

Chowfin, S.M. (2025). A Perspective on Twenty Years of Human-Leopard Conflict in Pauri Garhwal, Uttarakhand, India with Sustainable Solutions for its Mitigation. biorxiv. doi: https://doi.org/10.1101/2025.08.15.669212.

Shrestha, P., Bishwakarma, K.L., Maharjan, R. and Sharma, D.K. (2025a). Claws in the capital: Human–leopard conflict hotspots and community perceptions in Kathmandu Valley, Nepal. Ecology and Evolution, [online] 15(12). doi:10.1002/ece3.72678.

human fatalities and injuries caused by large mammals. PLOS ONE, [online] 11(9), p.e0161717. doi:10.1371/journal.pone.0161717.

Khorozyan, I., Ghoddousi, A., Soofi, M. and Waltert, M. (2015). Big cats kill more livestock when wild prey reaches a minimum threshold. Biological Conservation, 192, pp.268–275. doi:10.1016/j.biocon.2015.09.031.

attitudes towards the common leopardPanthera pardusin a protected area in the Indian himalayan region. Oryx, 58(2), pp.202–209. doi:10.1017/s0030605323001278.

(2022). Potential risk zone for anthropogenic mortality of carnivores in Gandaki Province, Nepal. Ecology and Evolution, 12(1). doi:10.1002/ece3.8491.

practices and herd composition influence leopard-human conflict in Pokhara Valley, Nepal. Human Dimensions of Wildlife, 25(1), pp.62–69. doi:10.1080/10871209.2019.1695157.

midhill landscape: A case study from panchase area in Chitwan Annapurna Landscape, Nepal. Journal of Institute of Science and Technology, [online] 23(1), pp.30–38. doi:10.3126/jist.v23i1.22158.

central part of Chitwan Annapurna Landscape, Nepal. Heliyon, [online] 10(4), p.e26386. doi:10.1016/j.heliyon.2024.e26386.

(2021). Inhabiting terra incognita: Two-decadal patterns of negative human-leopard interactions in human-dominating landscape of Maharashtra, India. Global Ecology and Conservation, 29, p.e01740.

leopardPanthera pardusdiet in a human-dominated landscape in western Maharashtra, India. Oryx, [online] 50(1), pp.156–162. doi:10.1017/s0030605314000106.

leopards in Shivapuri Nagarjun National Park, Nepal. Russian Journal of Ecology, 50(6), pp.590–592. doi:10.1134/s1067413619060031.

Assessing determinants of presence and predation on domestic animals. Oryx, [online] 56(1), pp.91–100. doi:10.1017/S0030605320000423.

Kala, C.P. and Kothari, K.K. (2013). Human–wildlife conflicts and conservation issues. HumanWildlife Interactions, [online] 7(2), pp.325–333. doi:10.2307/24874875.

of human-leopard conflict in the human-dominated landscape of mid-hills: A case study from Tanahun District of Nepal. International Journal of Zoology, 2023, pp.1–12. doi:10.1155/2023/5690289.

and compensation: Insights from western ghats protected areas. Biological Conservation, 166, pp.175–185. doi:10.1016/j.biocon.2013.06.027.

Lamichhane , S., Thapa, A., Thapa, M., Panthi, S. and Giarondo, A. (2022). Assessing human-leopard conflict in the middle mountain of western Nepal. Authorea. doi:https://doi.org/10.22541/au.164758652.25506348/v1.

and Adhikari, H. (2023). Landscape predictors influencing livestock depredation by leopards in and around Annapurna Conservation Area, Nepal. PeerJ, 11, p.e16516. doi:10.7717/peerj.16516.

Municipality, Lalitpur, Nepal. Journal of Institute of Science and Technology, 27(2), pp.1–11. doi:10.3126/jist.v27i2.51171.

leopard conservation in Terai arc landscape, India. Human-Wildlife Interactions, [online] 9(1), pp.48–57. Available at: https://www.jstor.org/stable/24874924 [Accessed 24 Sept. 2026].

leopards, associated economic losses, and financial compensation to communities in Chhota Udepur district of central Gujarat, India. Human-Wildlife Interactions, 17(1), pp.1–10. doi:https://doi.org/10.26077/5242-8408.

Trends of human-wildlife conflicting in Kali Tiger Reserve Forest, Karnataka, South India. International Journal of Environment and Climate Change, 13(12), pp.724–729. doi:10.9734/ijecc/2023/v13i123734.

municipality, Gorkha district, Nepal. Journal of Agriculture and Natural Resources, [online] 8(1), pp.20–29. doi:10.3126/janr.v8i1.88829.

Pun, P., Pramila Neupane, Tulasi Prasad Dahal, Trishna Rayamajhi, Prashamsa Paudel, Thapa, A., Pramod Raj Regmi, Shankar Thami, Thapa, G., Suraj Khanal, Lama, S., Karki, J., Sujan Khanal and Ferdin, A.E.J.

(2024). Human–wildlife conflict in Bardia—Banke complex: Patterns of human fatalities and injuries caused by large mammals. Ecology and Evolution, [online] 14(10). doi:10.1002/ece3.70395.

Ramesh, T., Kalle, R., Milda, D., Gayathri, V., Thanikodi, M., Ashish, K. and Giordano, A.J. (2020). Patterns of livestock predation risk by large carnivores in India’s eastern and western ghats. Global Ecology and Conservation, 24, p.e01366. doi:10.1016/j.gecco.2020.e01366.

habitat transformations on human-large carnivore interactions outside protected areas. Scientific Reports, [online] 15(1). doi:10.1038/s41598-025-13808-4.

Sharma, P. and Chishty, N. (2018). Human-panther conflicts in the Aravalli Hills of southern rajasthan-A case study. Indian Journal of Ecology, 45(3), pp.489–493. doi:10.13140/RG.2.2.13549.20967.

Wangdi, S., Dorji, T., Wangchuk, N., Chitale, V.S., Uprety, Y. and Sharma, E. (2020). Mapping human–wildlife conflict hotspots in a transboundary landscape, Eastern Himalaya. Global Ecology and Conservation, 24,

landscape in Anamalai Hills, India. Conservation and Society, [online] 15(4), pp.474–482. Available at: https://www.jstor.org/stable/26393316 [Accessed 24 Sept. 2026].

Sijapati, R.K., Sharma, H.P., Sharma, S., Subedi, J.R. and Belant, J.L. (2021). Livestock depredation by leopards and Tigers near Bardia National Park, Nepal. Animals, 11(7), p.1896. doi:10.3390/ani11071896.

economic losses in and around Hirpora Wildlife Sanctuary along Pir Panjal Moutain range, Western Himalayas. Journal of Wildlife and Biodiversity, [online] 6(Suppl. 1), pp.119–136. doi:10.5281/zenodo.6959846.

large carnivore conflict in the Chitwan National Park, Nepal. Journal of Resources and Ecology, 15(4). doi:10.5814/j.issn.1674-764x.2024.04.005.

Wang, S.W and Macdonald, D.W (2006). Livestock predation by carnivores in Jigme Singye Wangchuck National Park, Bhutan. Biological Conservation, [online] 129(4), pp.558–565. doi:10.1016/j.biocon.2005.11.024.

Park and Sanctuary, Gujarat, India. International Journal of Ecology and Environmental Sciences, 48(6). doi:10.55863/ijees.2022.6755.

Sathyamangalam and mudumalai tiger reserves. Tropical Conservation Science, 11, p.194008291880275. doi:10.1177/1940082918802758.
