## Supplementary material for "Understanding Spatial Patterns of Human-Tiger and Human-Leopard Conflict across South Asia using a Multiscale Regional Approach": S4

### Table S4. Predictive performance across all model configurations.

Human–leopard and human–tiger conflict models evaluated across predictor scales, spatial cross-validation block sizes and background-sampling methods.

| Species | Method | Predictor scale (km) | CV block size (km) | Model | PR-AUC, mean (SD) | ROC-AUC, mean (SD) | Brier score, mean (SD) | Log-loss, mean (SD) | Successful folds | Final predictors |
|---|---|---|---|---|---|---|---|---|---|---|
| HLC | AA | 1 | 10 | Full | 0.241 (0.041) | 0.722 (0.035) | 0.082 (0.005) | 0.292 (0.011) | 5 | 24 |
| HLC | AA | 1 | 20 | Full | 0.195 (0.046) | 0.689 (0.055) | 0.085 (0.005) | 0.303 (0.019) | 5 | 24 |
| HLC | AA | 1 | 30 | Full | 0.210 (0.120) | 0.698 (0.065) | 0.085 (0.007) | 0.299 (0.016) | 5 | 24 |
| HLC | AA | 1 | 40 | Full | 0.179 (0.060) | 0.675 (0.056) | 0.087 (0.021) | 0.311 (0.068) | 5 | 24 |
| HLC | AA | 1 | 50 | Full | 0.180 (0.071) | 0.674 (0.061) | 0.087 (0.008) | 0.308 (0.025) | 5 | 24 |
| HLC | AA | 2 | 10 | Full | 0.266 (0.061) | 0.733 (0.037) | 0.081 (0.006) | 0.289 (0.014) | 5 | 25 |
| HLC | AA | 2 | 20 | Full | 0.216 (0.048) | 0.699 (0.046) | 0.084 (0.005) | 0.300 (0.019) | 5 | 25 |
| HLC | AA | 2 | 30 | Full | 0.234 (0.141) | 0.710 (0.062) | 0.084 (0.007) | 0.297 (0.016) | 5 | 25 |
| HLC | AA | 2 | 40 | Full | 0.194 (0.080) | 0.685 (0.065) | 0.086 (0.021) | 0.310 (0.068) | 5 | 25 |
| HLC | AA | 2 | 50 | Full | 0.205 (0.108) | 0.686 (0.071) | 0.086 (0.007) | 0.307 (0.024) | 5 | 25 |
| HLC | AA | 5 | 10 | Full | 0.288 (0.064) | 0.748 (0.031) | 0.080 (0.006) | 0.285 (0.016) | 5 | 24 |
| HLC | AA | 5 | 20 | Full | 0.229 (0.041) | 0.713 (0.034) | 0.083 (0.005) | 0.298 (0.018) | 5 | 24 |
| HLC | AA | 5 | 30 | Full | 0.240 (0.135) | 0.720 (0.067) | 0.084 (0.007) | 0.295 (0.018) | 5 | 24 |
| HLC | AA | 5 | 40 | Full | 0.176 (0.041) | 0.686 (0.043) | 0.087 (0.020) | 0.313 (0.064) | 5 | 24 |
| HLC | AA | 5 | 50 | Full | 0.212 (0.131) | 0.685 (0.098) | 0.087 (0.007) | 0.311 (0.022) | 5 | 24 |
| HLC | BC | 1 | 10 | Full | 0.181 (0.030) | 0.677 (0.037) | 0.085 (0.005) | 0.303 (0.013) | 5 | 23 |

**Table S4. Predictive performance across all model configurations.**

Human–leopard and human–tiger conflict models evaluated across predictor scales, spatial cross-validation block sizes and background-sampling methods.

| Species | Method | Predictor scale (km) | CV block size (km) | Model | PR-AUC, mean (SD) | ROC-AUC, mean (SD) | Brier score, mean (SD) | Log-loss, mean (SD) | Successful folds | Final predictors |
|---|---|---|---|---|---|---|---|---|---|---|
| HLC | BC | 1 | 20 | Full | 0.166 (0.033) | 0.656 (0.057) | 0.086 (0.004) | 0.308 (0.012) | 5 | 23 |
| HLC | BC | 1 | 30 | Full | 0.151 (0.025) | 0.642 (0.055) | 0.087 (0.004) | 0.313 (0.018) | 5 | 23 |
| HLC | BC | 1 | 40 | Full | 0.160 (0.048) | 0.634 (0.095) | 0.088 (0.007) | 0.319 (0.029) | 5 | 23 |
| HLC | BC | 1 | 50 | Full | 0.135 (0.028) | 0.595 (0.060) | 0.090 (0.007) | 0.327 (0.026) | 5 | 23 |
| HLC | BC | 2 | 10 | Full | 0.200 (0.050) | 0.689 (0.043) | 0.085 (0.004) | 0.301 (0.013) | 5 | 24 |
| HLC | BC | 2 | 20 | Full | 0.185 (0.045) | 0.669 (0.063) | 0.086 (0.005) | 0.306 (0.015) | 5 | 24 |
| HLC | BC | 2 | 30 | Full | 0.167 (0.043) | 0.655 (0.067) | 0.087 (0.005) | 0.311 (0.020) | 5 | 24 |
| HLC | BC | 2 | 40 | Full | 0.182 (0.083) | 0.650 (0.110) | 0.087 (0.007) | 0.317 (0.033) | 5 | 24 |
| HLC | BC | 2 | 50 | Full | 0.137 (0.027) | 0.600 (0.049) | 0.090 (0.007) | 0.328 (0.022) | 5 | 24 |
| HLC | BC | 5 | 10 | Full | 0.222 (0.069) | 0.713 (0.044) | 0.083 (0.004) | 0.296 (0.012) | 5 | 22 |
| HLC | BC | 5 | 20 | Full | 0.201 (0.062) | 0.681 (0.068) | 0.085 (0.006) | 0.305 (0.019) | 5 | 22 |
| HLC | BC | 5 | 30 | Full | 0.181 (0.045) | 0.675 (0.071) | 0.086 (0.004) | 0.309 (0.020) | 5 | 22 |
| HLC | BC | 5 | 40 | Full | 0.179 (0.074) | 0.665 (0.112) | 0.088 (0.008) | 0.317 (0.036) | 5 | 22 |
| HLC | BC | 5 | 50 | Full | 0.132 (0.019) | 0.587 (0.073) | 0.091 (0.007) | 0.332 (0.022) | 5 | 22 |
| HLC | UR | 1 | 10 | Full | 0.211 (0.057) | 0.707 (0.054) | 0.084 (0.009) | 0.297 (0.033) | 5 | 23 |
| HLC | UR | 1 | 20 | Full | 0.179 (0.031) | 0.678 (0.050) | 0.086 (0.007) | 0.305 (0.023) | 5 | 23 |

### Table S4. Predictive performance across all model configurations.

Human–leopard and human–tiger conflict models evaluated across predictor scales, spatial cross-validation block sizes and background-sampling methods.

| Species | Method | Predictor scale (km) | CV block size (km) | Model | PR-AUC, mean (SD) | ROC-AUC, mean (SD) | Brier score, mean (SD) | Log-loss, mean (SD) | Successful folds | Final predictors |
|---|---|---|---|---|---|---|---|---|---|---|
| HLC | UR | 1 | 30 | Full | 0.170 (0.048) | 0.673 (0.073) | 0.087 (0.010) | 0.308 (0.031) | 5 | 23 |
| HLC | UR | 1 | 40 | Full | 0.167 (0.083) | 0.638 (0.113) | 0.088 (0.013) | 0.318 (0.043) | 5 | 23 |
| HLC | UR | 1 | 50 | Full | 0.132 (0.018) | 0.617 (0.045) | 0.090 (0.010) | 0.324 (0.033) | 5 | 23 |
| HLC | UR | 2 | 10 | Full | 0.228 (0.078) | 0.707 (0.053) | 0.083 (0.009) | 0.297 (0.033) | 5 | 23 |
| HLC | UR | 2 | 20 | Full | 0.187 (0.035) | 0.681 (0.052) | 0.086 (0.007) | 0.305 (0.023) | 5 | 23 |
| HLC | UR | 2 | 30 | Full | 0.181 (0.058) | 0.673 (0.067) | 0.087 (0.009) | 0.309 (0.028) | 5 | 23 |
| HLC | UR | 2 | 40 | Full | 0.176 (0.094) | 0.652 (0.115) | 0.088 (0.013) | 0.318 (0.047) | 5 | 23 |
| HLC | UR | 2 | 50 | Full | 0.139 (0.028) | 0.623 (0.045) | 0.090 (0.009) | 0.324 (0.030) | 5 | 23 |
| HLC | UR | 5 | 10 | Full | 0.248 (0.078) | 0.733 (0.049) | 0.082 (0.010) | 0.290 (0.035) | 5 | 23 |
| HLC | UR | 5 | 20 | Full | 0.204 (0.030) | 0.700 (0.064) | 0.085 (0.009) | 0.301 (0.027) | 5 | 23 |
| HLC | UR | 5 | 30 | Full | 0.184 (0.057) | 0.648 (0.139) | 0.086 (0.008) | 0.308 (0.025) | 5 | 23 |
| HLC | UR | 5 | 40 | Full | 0.193 (0.105) | 0.674 (0.123) | 0.087 (0.013) | 0.314 (0.051) | 5 | 23 |
| HLC | UR | 5 | 50 | Full | 0.143 (0.027) | 0.610 (0.110) | 0.090 (0.009) | 0.324 (0.029) | 5 | 23 |
| HTC | AA | 1 | 10 | Full | 0.453 (0.021) | 0.833 (0.017) | 0.063 (0.005) | 0.230 (0.012) | 5 | 23 |
| HTC | AA | 1 | 20 | Full | 0.420 (0.113) | 0.819 (0.044) | 0.065 (0.003) | 0.237 (0.009) | 5 | 23 |
| HTC | AA | 1 | 30 | Full | 0.434 (0.064) | 0.822 (0.031) | 0.064 (0.012) | 0.235 (0.034) | 5 | 23 |

### Table S4. Predictive performance across all model configurations.

Human–leopard and human–tiger conflict models evaluated across predictor scales, spatial cross-validation block sizes and background-sampling methods.

| Species | Method | Predictor scale (km) | CV block size (km) | Model | PR-AUC, mean (SD) | ROC-AUC, mean (SD) | Brier score, mean (SD) | Log-loss, mean (SD) | Successful folds | Final predictors |
|---|---|---|---|---|---|---|---|---|---|---|
| HTC | AA | 1 | 40 | Full | 0.380 (0.115) | 0.806 (0.041) | 0.067 (0.016) | 0.244 (0.047) | 5 | 23 |
| HTC | AA | 1 | 50 | Full | 0.346 (0.162) | 0.765 (0.129) | 0.073 (0.026) | 0.284 (0.126) | 5 | 23 |
| HTC | AA | 2 | 10 | Full | 0.336 (0.185) | 0.723 (0.150) | 0.076 (0.020) | 1.485 (1.723) | 5 | 25 |
| HTC | AA | 2 | 20 | Full | 0.423 (0.092) | 0.829 (0.021) | 0.065 (0.003) | 0.234 (0.008) | 5 | 25 |
| HTC | AA | 2 | 30 | Full | 0.371 (0.192) | 0.765 (0.153) | 0.068 (0.012) | 0.759 (1.172) | 5 | 25 |
| HTC | AA | 2 | 40 | Full | 0.393 (0.113) | 0.812 (0.046) | 0.067 (0.019) | 0.244 (0.055) | 5 | 25 |
| HTC | AA | 2 | 50 | Full | 0.355 (0.166) | 0.772 (0.136) | 0.072 (0.026) | 0.284 (0.131) | 5 | 25 |
| HTC | AA | 5 | 10 | Full | 0.491 (0.037) | 0.853 (0.020) | 0.061 (0.003) | 0.221 (0.006) | 5 | 26 |
| HTC | AA | 5 | 20 | Full | 0.444 (0.132) | 0.841 (0.027) | 0.063 (0.002) | 0.229 (0.004) | 5 | 26 |
| HTC | AA | 5 | 30 | Full | 0.449 (0.075) | 0.839 (0.026) | 0.063 (0.008) | 0.228 (0.020) | 5 | 26 |
| HTC | AA | 5 | 40 | Full | 0.421 (0.117) | 0.825 (0.046) | 0.065 (0.017) | 0.237 (0.046) | 5 | 26 |
| HTC | AA | 5 | 50 | Full | 0.338 (0.186) | 0.766 (0.147) | 0.074 (0.027) | 0.297 (0.152) | 5 | 26 |
| HTC | BC | 1 | 10 | Full | 0.424 (0.052) | 0.830 (0.014) | 0.065 (0.006) | 0.235 (0.016) | 5 | 22 |
| HTC | BC | 1 | 20 | Full | 0.393 (0.113) | 0.818 (0.045) | 0.067 (0.009) | 0.242 (0.032) | 5 | 22 |
| HTC | BC | 1 | 30 | Full | 0.384 (0.089) | 0.808 (0.041) | 0.067 (0.005) | 0.244 (0.019) | 5 | 22 |
| HTC | BC | 1 | 40 | Full | 0.310 (0.185) | 0.774 (0.093) | 0.073 (0.017) | 0.264 (0.055) | 5 | 22 |

### Table S4. Predictive performance across all model configurations.

Human–leopard and human–tiger conflict models evaluated across predictor scales, spatial cross-validation block sizes and background-sampling methods.

| Species | Method | Predictor scale (km) | CV block size (km) | Model | PR-AUC, mean (SD) | ROC-AUC, mean (SD) | Brier score, mean (SD) | Log-loss, mean (SD) | Successful folds | Final predictors |
|---|---|---|---|---|---|---|---|---|---|---|
| HTC | BC | 1 | 50 | Full | 0.294 (0.222) | 0.729 (0.152) | 0.073 (0.024) | 0.274 (0.090) | 5 | 22 |
| HTC | BC | 2 | 10 | Full | 0.447 (0.045) | 0.841 (0.009) | 0.064 (0.005) | 0.230 (0.012) | 5 | 25 |
| HTC | BC | 2 | 20 | Full | 0.384 (0.103) | 0.820 (0.049) | 0.068 (0.009) | 0.243 (0.033) | 5 | 25 |
| HTC | BC | 2 | 30 | Full | 0.400 (0.118) | 0.816 (0.047) | 0.066 (0.006) | 0.241 (0.022) | 5 | 25 |
| HTC | BC | 2 | 40 | Full | 0.295 (0.163) | 0.773 (0.091) | 0.073 (0.015) | 0.265 (0.049) | 5 | 25 |
| HTC | BC | 2 | 50 | Full | 0.299 (0.200) | 0.748 (0.136) | 0.073 (0.022) | 0.279 (0.094) | 5 | 25 |
| HTC | BC | 5 | 10 | Full | 0.477 (0.068) | 0.854 (0.010) | 0.061 (0.005) | 0.222 (0.012) | 5 | 25 |
| HTC | BC | 5 | 20 | Full | 0.417 (0.099) | 0.836 (0.041) | 0.066 (0.008) | 0.235 (0.030) | 5 | 25 |
| HTC | BC | 5 | 30 | Full | 0.434 (0.097) | 0.828 (0.043) | 0.064 (0.005) | 0.234 (0.017) | 5 | 25 |
| HTC | BC | 5 | 40 | Full | 0.312 (0.166) | 0.784 (0.085) | 0.072 (0.016) | 0.262 (0.053) | 5 | 25 |
| HTC | BC | 5 | 50 | Full | 0.318 (0.198) | 0.757 (0.129) | 0.072 (0.022) | 0.280 (0.100) | 5 | 25 |
| HTC | UR | 1 | 10 | Full | 0.417 (0.067) | 0.831 (0.013) | 0.065 (0.002) | 0.234 (0.005) | 5 | 22 |
| HTC | UR | 1 | 20 | Full | 0.435 (0.065) | 0.835 (0.031) | 0.065 (0.013) | 0.235 (0.041) | 5 | 22 |
| HTC | UR | 1 | 30 | Full | 0.320 (0.097) | 0.796 (0.041) | 0.072 (0.015) | 0.259 (0.044) | 5 | 22 |
| HTC | UR | 1 | 40 | Full | 0.351 (0.105) | 0.805 (0.027) | 0.070 (0.007) | 0.251 (0.017) | 5 | 22 |
| HTC | UR | 1 | 50 | Full | 0.293 (0.206) | 0.739 (0.123) | 0.075 (0.024) | 0.284 (0.092) | 5 | 22 |

### Table S4. Predictive performance across all model configurations.

Human–leopard and human–tiger conflict models evaluated across predictor scales, spatial cross-validation block sizes and background-sampling methods.

| Species | Method | Predictor scale (km) | CV block size (km) | Model | PR-AUC, mean (SD) | ROC-AUC, mean (SD) | Brier score, mean (SD) | Log-loss, mean (SD) | Successful folds | Final predictors |
|---|---|---|---|---|---|---|---|---|---|---|
| HTC | UR | 2 | 10 | Full | 0.442 (0.069) | 0.842 (0.015) | 0.064 (0.003) | 0.229 (0.009) | 5 | 25 |
| HTC | UR | 2 | 20 | Full | 0.445 (0.073) | 0.840 (0.032) | 0.065 (0.013) | 0.233 (0.042) | 5 | 25 |
| HTC | UR | 2 | 30 | Full | 0.330 (0.081) | 0.803 (0.043) | 0.071 (0.015) | 0.258 (0.046) | 5 | 25 |
| HTC | UR | 2 | 40 | Full | 0.398 (0.103) | 0.821 (0.029) | 0.067 (0.006) | 0.243 (0.018) | 5 | 25 |
| HTC | UR | 2 | 50 | Full | 0.309 (0.217) | 0.748 (0.127) | 0.074 (0.022) | 0.282 (0.095) | 5 | 25 |
| HTC | UR | 5 | 10 | Full | 0.485 (0.065) | 0.856 (0.014) | 0.061 (0.003) | 0.221 (0.006) | 5 | 26 |
| HTC | UR | 5 | 20 | Full | 0.466 (0.037) | 0.852 (0.019) | 0.063 (0.011) | 0.227 (0.034) | 5 | 26 |
| HTC | UR | 5 | 30 | Full | 0.371 (0.065) | 0.823 (0.030) | 0.069 (0.013) | 0.248 (0.039) | 5 | 26 |
| HTC | UR | 5 | 40 | Full | 0.420 (0.072) | 0.831 (0.021) | 0.065 (0.007) | 0.238 (0.019) | 5 | 26 |
| HTC | UR | 5 | 50 | Full | 0.330 (0.204) | 0.762 (0.128) | 0.072 (0.023) | 0.279 (0.102) | 5 | 26 |

Notes: HLC, human–leopard conflict; HTC, human–tiger conflict; AA, Accessible Area; BC, Bias Corrected; UR, Uniform Random; CV, cross-validation; SD, standard deviation.

PR-AUC is the area under the precision–recall curve; ROC-AUC is the area under the receiver operating characteristic curve. Higher values indicate better discrimination.

Lower Brier scores and log-loss indicate better probabilistic predictive performance. Values are reported as the saved mean followed by the saved SD in parentheses.

SDs describe variation among the evaluations summarized in the source files and are not confidence intervals. Successful folds and final predictor counts are reported as saved.

Every saved model configuration is included. Values are displayed to three decimal places; no additional averaging across background-sampling methods was performed.
