## Supplementary Materials and Results for "Understanding Spatial Patterns of Human-Tiger and Human-Leopard Conflict across South Asia using a Multiscale Regional Approach"

### Supplementary methods and results

#### S1. Predictor variable extraction

Predictors represented landscape structure, anthropogenic disturbance, bioclimatic conditions, topography and hydrology. Unless stated otherwise, spatial summaries were calculated within 1, 2 and 5km circular buffers around conflict and background locations. Table A1 lists variable definitions, sources, native resolutions, extraction extents and temporal matching.

##### S1.1. Landscape structure

Landscape composition and configuration may influence conflict by shaping carnivore distribution, movement, prey availability and exposure to human activity (Fahrig, 2007; McIntyre and Wiens, 1999). Forest cover, fragmentation, habitat loss and heterogeneity have also been associated with conflict risk (Acharya *et al*., 2017; Haddad *et al*., 2015; Joshi *et al*., 2016; Lamichhane *et al*., 2023; Miller, Jhala and Schmitz, 2016; Naha *et al*., 2020). Landscape structure was characterized using land-cover composition, Shannon's Diversity Index (SHDI), forest patch density, edge density and mean patch area, distance to forest, baseline tree-canopy cover, cumulative tree-cover loss and NDVI.

Land-cover composition and forest configuration were derived from the 30-m Global Land Analysis and Discovery (GLAD) land-cover products (Potapov *et al*., 2022). Records from 2000–2004, 2005–2009, 2010–2014, 2015–2019 and 2020 onwards were matched to the 2000, 2005, 2010, 2015 and 2020 products, respectively. GLAD classes were consolidated into cropland (1–3), forest (5–13), shrubland (14–16 and 21), grassland (17), built-up land (26), bare ground (27–29), water (30) and other classes. Category cover was calculated as the mean of a binary class mask multiplied by 100. Landscape heterogeneity was quantified as SHDI = $-\Sigma(p_i \ln p_i)$, where $p_i$ is the proportion occupied by category i (McGarigal, Cushman and Ene, 2012).

Forest configuration was calculated from temporally matched forest masks using *landscapemetrics* in R (Hesselbarth *et al*., 2019). Forest cells sharing an edge or corner formed a patch. Patch density, edge density and mean patch area were expressed as patches per 100 ha, m $ha^{-1}$ and hectares, respectively. Euclidean distance to the nearest forest pixel was calculated separately for each GLAD year at 30-m resolution and extracted at each location in kilometres.

Baseline canopy cover and cumulative loss were derived from Hansen Global Forest Change version 1.12 at 30-m resolution (Hansen *et al*., 2013). Summaries included mean canopy cover in 2000, cumulative loss by the observation year as a percentage of buffer area, and cumulative loss relative to canopy-weighted baseline cover. NDVI was derived from cloud-, shadow- and snow-screened Landsat Collection 2 Level-2 surface reflectance at 30-m resolution (Vermote *et al*., 2016). Annual median composites were matched to observation years and mean NDVI was calculated within each buffer (Tucker, 1979).

##### S1.2. Anthropogenic disturbance

Anthropogenic activity may affect conflict through its influence on carnivore occurrence, movement and overlap with people and livestock (Naha *et al*., 2020; Rostro-García *et al*., 2016; Sunarto *et al*., 2012). Human population density, built-up cover and distance to settlements represented human presence; road density and distance to roads represented infrastructure and accessibility; and livestock density represented potential domestic-prey availability (Carter *et al*., 2020; Lamichhane *et al*., 2023b; Naha *et al*., 2020; Quintana *et al*., 2022). Built-up cover was calculated from the GLAD categories described above.

Human population density was extracted at each location from Gridded Population of the World version 4 at 30-arc-second resolution and expressed as people $km^{-2}$ (CIESIN, 2017). Records from 2000–2004, 2005–2009, 2010–2014, 2015–2019 and 2020 onwards were matched to the 2000, 2005, 2010, 2015 and 2020 rasters, respectively.

Distance to settlements was derived from the 100-m GHS-BUILT-S R2023A product (Pesaresi and Politis, 2026). Pixels with built-up values above zero were classified as settlements. Euclidean distance to the nearest such pixel was calculated in kilometres at 100-m resolution, with a maximum search radius of 50 km. Records from 2000–2007 were assigned the 2000 layer and records from 2008 onwards the 2015 layer.

Road variables were derived from the Global Roads Inventory Project dataset (Meijer *et al*., 2018). Road density was the total mapped road length divided by buffer area (km km$^{-2}$); distance to roads was the shortest Euclidean distance to a mapped road (km). The single road-network layer was treated as temporally static.

Livestock density was obtained from the FAO Gridded Livestock of the World products for 2010, 2015 and 2020 (FAO, 2024, 2026; Gilbert *et al*., 2018). Dasymetric products allocate census counts using environmental and land-use covariates. Species-specific densities for buffalo, cattle, chickens, goats, pigs and sheep were summed for 2010 and 2015; the combined 2020 product was used directly. Records through 2010 were matched to 2010, records from 2011–2019 to 2015, and records from 2020 onwards to 2020. Livestock density was averaged within a 1km circular buffer and retained in the units of the source density raster.

##### S1.3. Bioclimatic variables

Temperature and precipitation may influence resource availability, carnivore distributions and overlap with people and livestock (Calhoun *et al*., 2025; Lamichhane *et al*., 2023b; Sharma *et al*., 2020). The supplied candidate-variable inventory included isothermality (BIO3), maximum temperature of the warmest month (BIO5), minimum temperature of the coldest month (BIO6), annual precipitation (BIO12), and precipitation of the wettest and driest quarters (BIO16 and BIO17). These variables were obtained from WorldClim version 2.1 at 30-arc-second resolution and extracted at each location using *terra* in R (Fick and Hijmans, 2017). WorldClim represents long-term climatic averages, so layers were not matched to individual observation years.

##### S1.4. Topography

Topography may affect carnivore habitat use and the accessibility of terrain to people and livestock (Baral *et al*., 2023; Karki *et al*., 2025; Sharma *et al*., 2020). Elevation, slope and aspect were derived in Google Earth Engine from the 30-m Copernicus DEM GLO-30 product (European Space Agency, 2024). Mean elevation and slope were calculated within each buffer. Aspect was summarized using its circular mean and transformed to northness (cosine) and eastness (sine) (Amatulli *et al*., 2018).

##### S1.5. Hydrology

Surface water may influence conflict by affecting carnivore movement and the distribution of wild prey, livestock and people (Miller, Jhala and Jena, 2015; Ramesh *et al*., 2020; Silwal *et al*., 2024). Water occurrence, water seasonality and distance to water were obtained from Joint Research Centre Global Surface Water version 1.4 at 30-m resolution (Pekel *et al*., 2016). Spatial means of the fixed occurrence and seasonality layers were calculated in each buffer; these were not matched to observation year. Occurrence represented the frequency of water detection over 1984–2021. Seasonality was the spatial mean of the source seasonality band, expressed as months of water presence; this spatial averaging does not constitute a temporal average across years. Masked pixels were set to zero before spatial averaging. Distance to the nearest pixel with occurrence greater than zero was calculated at nominal 30-m resolution and expressed in kilometres.

#### Table A1. Predictor definitions and extraction procedures

The table distinguishes point extraction from buffer summaries. Definitions refer to unstandardized predictors; continuous variables were subsequently standardized using training-fold parameters for cross-validation.

| Predictor | Definition / extraction | Units | Data source and native resolution | Spatial extent | Temporal matching |
|---|---|---|---|---|---|
| Cropland cover (Landscape) | The percentage of the surrounding circular extent occupied by cropland was calculated as the mean of a binary mask multiplied by 100. The supplied extraction code used land-cover classes 1–3. | % | GLAD land-cover rasters (Potapov et al., 2022) Native resolution: 30 m | Calculated within 1, 2 and 5km circular extents. | Records from 2000–2004, 2005–2009, 2010–2014 and 2015–2019 were matched to the 2000, 2005, 2010 and 2015 layers, respectively. Records from 2020 onwards were matched to the 2020 layer. |
| Forest cover (Landscape) | The percentage of the surrounding circular extent occupied by forest was calculated as the mean of a binary mask multiplied by 100. The supplied extraction code used land-cover classes 5–13. | % | GLAD land-cover rasters (Potapov et al., 2022) Native resolution: 30 m | Calculated within 1, 2 and 5km circular extents. | Records from 2000–2004, 2005–2009, 2010–2014 and 2015–2019 were matched to the 2000, 2005, 2010 and 2015 layers, respectively. Records from 2020 onwards were matched to the 2020 layer. |
| Shrubland cover (Landscape) | The percentage of the surrounding circular extent occupied by shrubland was calculated as the mean of a binary mask multiplied by 100. The supplied extraction code used land-cover classes 14–16 and 21. | % | GLAD land-cover rasters (Potapov et al., 2022) Native resolution: 30 m | Calculated within 1, 2 and 5km circular extents. | Records from 2000–2004, 2005–2009, 2010–2014 and 2015–2019 were matched to the 2000, 2005, 2010 and 2015 layers, respectively. Records from 2020 onwards were matched to the 2020 layer. |
| Grassland cover (Landscape) | The percentage of the surrounding circular extent occupied by grassland was calculated as the mean of a binary mask multiplied by 100. The supplied extraction code used land-cover classes 17. | % | GLAD land-cover rasters (Potapov et al., 2022) Native resolution: 30 m | Calculated within 1, 2 and 5km circular extents. | Records from 2000–2004, 2005–2009, 2010–2014 and 2015–2019 were matched to the 2000, 2005, 2010 and 2015 layers, respectively. Records from 2020 onwards were matched to the 2020 layer. |
| Bare-ground cover (Landscape) | The percentage of the surrounding circular extent occupied by bare-ground was calculated as the mean of a binary mask multiplied by 100. The supplied extraction code used land-cover classes 27–29. | % | GLAD land-cover rasters (Potapov et al., 2022) Native resolution: 30 m | Calculated within 1, 2 and 5km circular extents. | Records from 2000–2004, 2005–2009, 2010–2014 and 2015–2019 were matched to the 2000, 2005, 2010 and 2015 layers, respectively. Records from 2020 onwards were matched to the 2020 layer. |
| Shannon diversity index (Landscape) | Landscape diversity was calculated as $-\Sigma(p_i \ln p_i)$, where $p_i$ represents the area proportion of each consolidated land-cover class. Classes with zero area contributed zero. | Unitless | GLAD land-cover rasters (Potapov et al., 2022) Native resolution: 30 m | Calculated within 1, 2 and 5km circular extents. | Records from 2000–2004, 2005–2009, 2010–2014 and 2015–2019 were matched to the 2000, 2005, 2010 and 2015 layers, respectively. Records from 2020 onwards were matched to the 2020 layer. |
| Forest patch density (Landscape) | The number of forest patches was divided by the area of the surrounding landscape and expressed per 100 ha. Forest pixels sharing an edge or corner were treated as connected. | Patches per 100 ha | GLAD land-cover rasters (Potapov et al., 2022) Native resolution: 30 m | Calculated within 1, 2 and 5km circular extents. | Records from 2000–2004, 2005–2009, 2010–2014 and 2015–2019 were matched to the 2000, 2005, 2010 and 2015 layers, respectively. Records from 2020 onwards were matched to the 2020 layer. |

| Predictor | Definition / extraction | Units | Data source and native resolution | Spatial extent | Temporal matching |
|---|---|---|---|---|---|
| Forest edge density (Landscape) | Forest edge length was divided by the area of the surrounding landscape using the class-level edge-density metric in landscapemetrics. | m/ha | GLAD land-cover rasters (Potapov et al., 2022)<br>Native resolution: 30 m | Calculated within 1, 2 and 5km circular extents. | Records from 2000–2004, 2005–2009, 2010–2014 and 2015–2019 were matched to the 2000, 2005, 2010 and 2015 layers, respectively. Records from 2020 onwards were matched to the 2020 layer. |
| Mean forest patch area (Landscape) | Mean forest patch area was calculated using eight-neighbour connectivity, with forest pixels sharing an edge or corner treated as connected. | ha | GLAD land-cover rasters (Potapov et al., 2022)<br>Native resolution: 30 m | Calculated within 1, 2 and 5km circular extents. | Records from 2000–2004, 2005–2009, 2010–2014 and 2015–2019 were matched to the 2000, 2005, 2010 and 2015 layers, respectively. Records from 2020 onwards were matched to the 2020 layer. |
| Distance to forest (Landscape) | Euclidean distance to the nearest pixel assigned to forest classes 5–13 was calculated using a distance transform and extracted at each location. | km | GLAD land-cover rasters (Potapov et al., 2022)<br>Native resolution: 30 m | Extracted at each location. | Records from 2000–2004, 2005–2009, 2010–2014 and 2015–2019 were matched to the 2000, 2005, 2010 and 2015 layers, respectively. Records from 2020 onwards were matched to the 2020 layer. |
| Tree-cover loss relative to landscape area (Landscape) | The summed area of pixels with recorded tree-cover loss was divided by the total area of the surrounding circular extent and multiplied by 100. | % | Hansen Global Forest Change v1.12 (Hansen et al., 2013)<br>Native resolution: 30 m | Calculated within 1, 2 and 5km circular extents. | Loss was accumulated from 2001 to the observation year. For observations after 2024, loss was accumulated through 2024. Values were set to missing for records before 2001. |
| Tree-cover loss relative to baseline (Landscape) | Baseline canopy-weighted area in pixels experiencing loss was divided by total baseline canopy-weighted area and multiplied by 100. Each pixel contributed its area multiplied by its 2000 canopy-cover fraction. No binary forest threshold was applied. | % | Hansen Global Forest Change v1.12 (Hansen et al., 2013)<br>Native resolution: 30 m | Calculated within 1, 2 and 5km circular extents. | Canopy cover in 2000 defined the baseline. Loss was accumulated from 2001 to the observation year, capped at 2024. Values were set to missing for records before 2001. |
| NDVI (Landscape) | NDVI was calculated from scaled red and near-infrared surface reflectance after cloud, shadow and snow screening. Annual median NDVI composites were averaged within each circular extent. | Unitless | Landsat Collection 2 Level-2 (Vermote et al., 2016; Tucker, 1979)<br>Native resolution: 30 m | Calculated within 1, 2 and 5km circular extents. | Imagery from 1 January to 31 December of each observation year was used. Landsat 5 and 7 were combined for 2000–2012, and Landsat 7 and 8 were combined from 2013 onwards. |
| Built-up cover (Anthropogenic) | The percentage of the surrounding circular extent occupied by built-up was calculated as the mean of a binary mask multiplied by 100. The supplied extraction code used land-cover classes 26. | % | GLAD land-cover rasters (Potapov et al., 2022)<br>Native resolution: 30 m | Calculated within 1, 2 and 5km circular extents. | Records from 2000–2004, 2005–2009, 2010–2014 and 2015–2019 were matched to the 2000, 2005, 2010 and 2015 layers, respectively. Records from 2020 onwards were matched to the 2020 layer. |

| Predictor | Definition / extraction | Units | Data source and native resolution | Spatial extent | Temporal matching |
|---|---|---|---|---|---|
| Human population density (Anthropogenic) | Human population density was extracted from the corresponding GPW raster at each location. | People/km2 | GPW v4 (CIESIN, 2017)<br>Native resolution: 30 arc-seconds (approximately 1 km at the equator) | Extracted at each location. | Records from 2000–2004, 2005–2009, 2010–2014 and 2015–2019 were matched to the 2000, 2005, 2010 and 2015 layers, respectively. Records from 2020 onwards were matched to the 2020 layer. |
| Distance to settlements (Anthropogenic) | Euclidean distance to the nearest pixel with built-up surface greater than zero was calculated at 100-m resolution using a maximum search radius of 50 km. | km | GHS-BUILT-S R2023A (Pesaresi and Politis, 2026)<br>Native resolution: 100 m | Extracted at each location. | Records from 2000–2007 were matched to the 2000 built-up layer. Records from 2008 onwards were matched to the 2015 layer. |
| Road density (Anthropogenic) | Total mapped road length within each circular extent was divided by the area of that extent. | km/km2 | GRIP4 (Meijer et al., 2018)<br>Native resolution: Vector network | Calculated within 1, 2 and 5km circular extents. | The same GRIP4 road network was used for all observation years. No temporal matching was performed. |
| Distance to roads (Anthropogenic) | The shortest Euclidean distance from each location to the nearest mapped road was calculated. | km | GRIP4 (Meijer et al., 2018)<br>Native resolution: Vector network | Extracted at each location. | The same GRIP4 road network was used for all observation years. No temporal matching was performed. |
| Livestock density (Anthropogenic) | Species-specific density rasters were summed for 2010 and 2015. The combined raster was used for 2020. Mean values were extracted within a 1km circular extent around each location. | Native raster density units | GLW (Gilbert et al., 2018; FAO, 2024, 2026)<br>Native resolution: 5 arc-minutes (approximately 10 km at the equator) | A fixed 1km circular extent was used. | Records up to and including 2010 were matched to the 2010 layer. Records from 2011–2019 were matched to the 2015 layer. Records from 2020 onwards were matched to the 2020 layer. |
| Temperature seasonality (BIO4) (Climate) | Temperature seasonality was represented by the standard deviation of monthly mean temperature multiplied by 100. | °C × 100 | WorldClim v2.1 (Fick and Hijmans, 2017)<br>Native resolution: 30 arc-seconds (approximately 1 km at the equator) | Extracted at each location. | The same WorldClim v2.1 climatic averages were used for all observation years. No annual temporal matching was performed. |
| Isothermality (BIO3) (Climate) | Isothermality was calculated as the mean diurnal temperature range divided by the annual temperature range and multiplied by 100. | Index | WorldClim v2.1 (Fick and Hijmans, 2017)<br>Native resolution: 30 arc-seconds (approximately 1 km at the equator) | Extracted at each location. | The same WorldClim v2.1 climatic averages were used for all observation years. No annual temporal matching was performed. |
| Maximum temperature (BIO5) (Climate) | The maximum temperature of the warmest month was extracted at each location. | Degrees C | WorldClim v2.1 (Fick and Hijmans, 2017)<br>Native resolution: 30 arc-seconds (approximately 1 km at the equator) | Extracted at each location. | The same WorldClim v2.1 climatic averages were used for all observation years. No annual temporal matching was performed. |

| Predictor | Definition / extraction | Units | Data source and native resolution | Spatial extent | Temporal matching |
|---|---|---|---|---|---|
| Minimum temperature (BIO6) (Climate) | The minimum temperature of the coldest month was extracted at each location. | Degrees C | WorldClim v2.1 (Fick and Hijmans, 2017) Native resolution: 30 arc-seconds (approximately 1 km at the equator) | Extracted at each location. | The same WorldClim v2.1 climatic averages were used for all observation years. No annual temporal matching was performed. |
| Annual precipitation (BIO12) (Climate) | Total annual precipitation was extracted at each location. | mm | WorldClim v2.1 (Fick and Hijmans, 2017) Native resolution: 30 arc-seconds (approximately 1 km at the equator) | Extracted at each location. | The same WorldClim v2.1 climatic averages were used for all observation years. No annual temporal matching was performed. |
| Wettest-quarter precipitation (BIO16) (Climate) | Total precipitation during the wettest three-month quarter was extracted at each location. | mm | WorldClim v2.1 (Fick and Hijmans, 2017) Native resolution: 30 arc-seconds (approximately 1 km at the equator) | Extracted at each location. | The same WorldClim v2.1 climatic averages were used for all observation years. No annual temporal matching was performed. |
| Driest-quarter precipitation (BIO17) (Climate) | Total precipitation during the driest three-month quarter was extracted at each location. | mm | WorldClim v2.1 (Fick and Hijmans, 2017) Native resolution: 30 arc-seconds (approximately 1 km at the equator) | Extracted at each location. | The same WorldClim v2.1 climatic averages were used for all observation years. No annual temporal matching was performed. |
| Elevation (Topography) | Mean elevation was calculated within each circular extent. | m | Copernicus DEM GLO-30 (ESA, 2024; Amatulli et al., 2018) Native resolution: 30 m | Calculated within 1, 2 and 5km circular extents. | The same Copernicus DEM was used for all observation years because terrain was treated as temporally static. |
| Slope (Topography) | Mean terrain slope was calculated within each circular extent. | Degrees | Copernicus DEM GLO-30 (ESA, 2024; Amatulli et al., 2018) Native resolution: 30 m | Calculated within 1, 2 and 5km circular extents. | The same Copernicus DEM was used for all observation years because terrain was treated as temporally static. |
| Northness (Topography) | Northness was calculated as the cosine of circular mean aspect after conversion to radians. Circular mean aspect was obtained from the mean sine and cosine of pixel-level aspect. | -1 to 1 | Copernicus DEM GLO-30 (ESA, 2024; Amatulli et al., 2018) Native resolution: 30 m | Calculated within 1, 2 and 5km circular extents. | The same Copernicus DEM was used for all observation years because terrain was treated as temporally static. |

| Predictor | Definition / extraction | Units | Data source and native resolution | Spatial extent | Temporal matching |
|---|---|---|---|---|---|
| Eastness (Topography) | Eastness was calculated as the sine of circular mean aspect after conversion to radians. Circular mean aspect was obtained from the mean sine and cosine of pixel-level aspect. | -1 to 1 | Copernicus DEM GLO-30 (ESA, 2024; Amatulli et al., 2018) Native resolution: 30 m | Calculated within 1, 2 and 5km circular extents. | The same Copernicus DEM was used for all observation years because terrain was treated as temporally static. |
| Distance to water (Hydrology) | Euclidean distance to the nearest surface-water pixel was calculated using a distance transform. Water pixels were defined by occurrence values greater than zero in JRC Global Surface Water v1.4. | km | JRC Global Surface Water v1.4 (Pekel et al., 2016) Native resolution: 30 m | Extracted at each location. | The water mask was derived from the 1984–2021 occurrence layer and used for all observation years. No temporal matching was performed. |
| Water occurrence (Hydrology) | Mean surface-water occurrence was calculated within each circular extent from the JRC occurrence band. Masked pixels were assigned zero before spatial averaging. | % | JRC Global Surface Water v1.4 (Pekel et al., 2016) Native resolution: 30 m | Calculated within 1, 2 and 5km circular extents. | The occurrence layer summarized water presence during 1984–2021 and was used for all observation years. No temporal matching was performed. |
| Water seasonality (Hydrology) | The number of months with surface water was averaged within each circular extent using the JRC seasonality band. Masked pixels were assigned zero before spatial averaging. | Months | JRC Global Surface Water v1.4 (Pekel et al., 2016) Native resolution: 30 m | Calculated within 1, 2 and 5km circular extents. | The same JRC Global Surface Water v1.4 seasonality layer was used for all observation years. No observation-year matching or averaging across annual layers was performed. |

#### Additional extraction details

Baseline canopy cover was calculated from Hansen treecover2000 as the spatial mean percentage canopy cover. Canopy-weighted baseline area was calculated by multiplying pixel area by the canopy-cover fraction. For baseline-relative loss, the same weighting was applied to pixels with lossyear greater than zero and no greater than the observation year minus 2000. The ratio was multiplied by 100. Where baseline canopy-weighted area was zero, the extraction code assigned a value of zero.

For observations after 2024, Hansen loss accumulation ended in 2024. The supplied code assigned missing values to all three Hansen outputs for records before 2001, including baseline canopy cover. Baseline canopy cover was an extracted quantity and an input to loss calculations but was not listed as a separate candidate in the supplied 33-variable consensus table. Temperature seasonality (BIO4) was included in that candidate list.

The JRC water-distance calculation used an occurrence threshold greater than zero and a distance-transform neighborhood of 2,048 pixels. Pixel distances were converted to kilometres using a nominal 30-m pixel size. Occurrence and seasonality were taken directly from the fixed v1.4 mapping layers. The code assigned zero to masked pixels before calculating spatial means and did not match hydrological layers to individual observation years.

The livestock script calculated mean values within 1km circular extents. Values were retained in the units of the input density rasters. Species-specific rasters were summed for 2010 and 2015, whereas the combined 2020 raster was used directly. The script did not apply a conversion between animal counts and livestock units.

Land-cover classes were consolidated into cropland (1–3), forest (5–13), shrubland (14–16 and 21), grassland (17), built-up land (26), bare ground (27–29), water (30) and other classes. Water and other cover proportions, forest aggregation and forest disturbance were among the quantities extracted by the supplied scripts but were not included in the supplied 33-variable HLC consensus inventory.

#### Supplementary results

##### S2. Spatial scale of conflict models

The predictive performance of binomial GLMs was compared across 45 configurations per species: three predictor scales (1, 2 and 5 km), five spatial cross-validation block sizes (10, 20, 30, 40 and 50 km) and three background-sampling methods. Models were evaluated using five-fold spatial cross-validation. PR-AUC was the primary model-selection criterion; ROC-AUC, Brier score and log-loss provided complementary measures. For each scale–block combination, PR-AUC was averaged first across folds within each background method, then across the three methods.

For HLC, mean PR-AUC generally increased with predictor scale and declined with increasing block size (Fig. S1A). The 5km scale outperformed the 1km scale at block sizes of 10, 20, 30 and 50 km. At 40 km, the 2km and 5km scales performed similarly (0.184 and 0.183, respectively). At the 5km predictor scale, mean PR-AUC declined from 0.253 at 10-km blocks to 0.211, 0.202, 0.183 and 0.162 at 20-, 30-, 40- and 50-km blocks, respectively.

For HTC, mean PR-AUC was highest at the 5km predictor scale at every block size (Fig. S1B). At this scale, it declined from 0.485 at 10-km blocks to 0.442, 0.418, 0.385 and 0.329 at 20-, 30-, 40- and 50-km blocks, respectively. The ordering of the 1 and 2km scales varied among block sizes. At the 2km scale, mean PR-AUC rose slightly between 10- and 20-km blocks before declining with larger blocks.

The combination of a 5km predictor scale and 10-km cross-validation blocks produced the highest mean PR-AUC for both HLC (0.253) and HTC (0.485) and was retained for subsequent analyses.

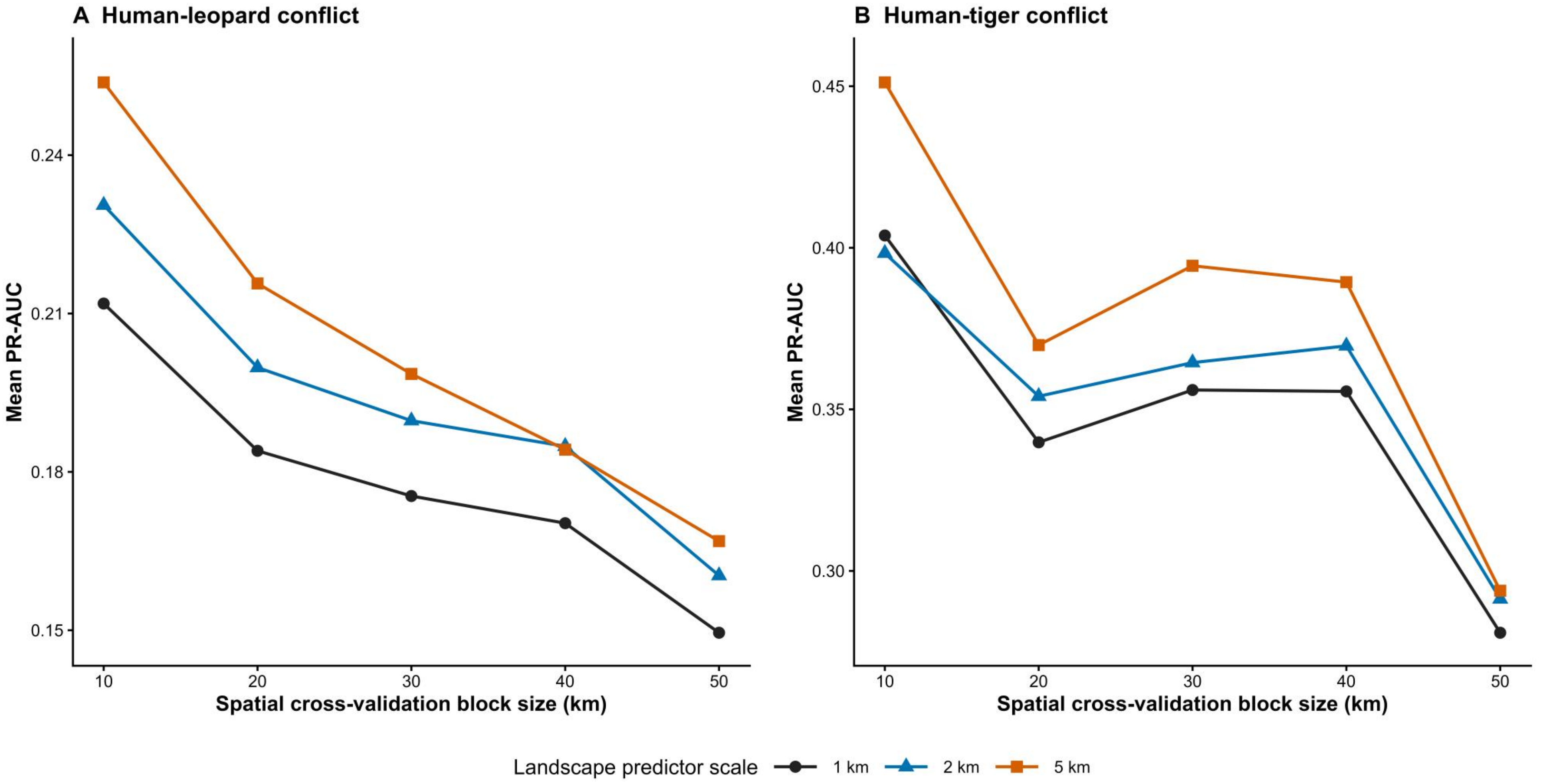


**Figure S1. Predictive performance of human–leopard and human–tiger conflict models was highest at the 5-km landscape predictor scale and 10-km spatial cross-validation block size.** Binomial GLMs for (A) human-leopard conflict (HLC) and (B) human-tiger conflict (HTC) were evaluated using five-fold spatial cross-validation across three landscape predictor scales (1, 2 and 5 km), five spatial block sizes (10, 20, 30, 40 and 50 km) and three background-sampling methods, yielding 45 model configurations per species. For each of the 15 predictor-scale and block-size combinations, PR-AUC was averaged across the three background-sampling methods, with performance within each method first averaged across the five spatial cross-validation folds. Predictor scales are distinguished by symbol and colour (1 km: circles; 2 km: triangles; 5 km: squares). For HLC, predictive performance generally increased with broader predictor scales and declined with increasing block size. For HTC, performance was highest at the 5-km predictor scale across all block sizes and declined progressively with increasing block size at this predictor scale.

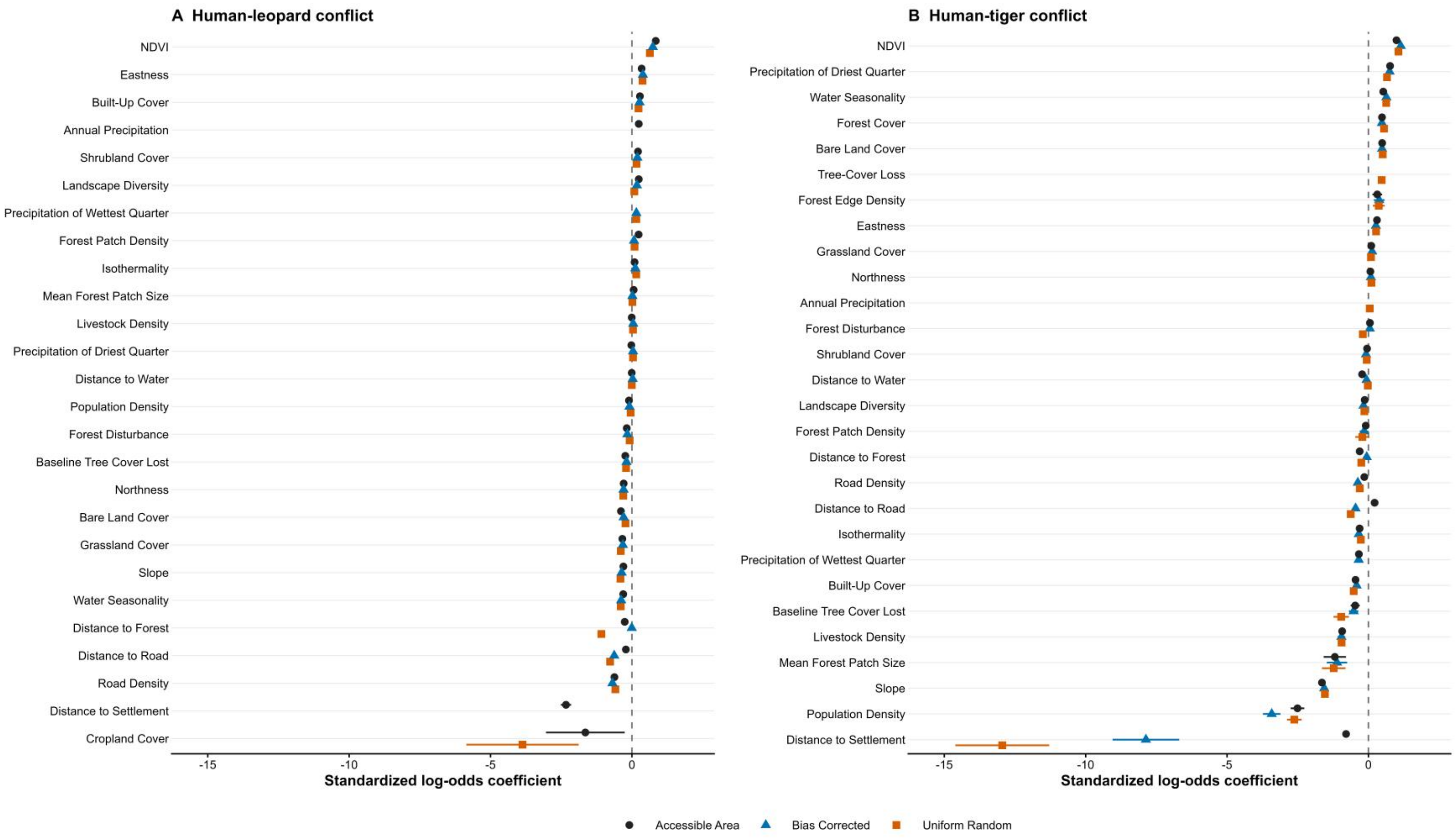


**Figure S2. Standardized model coefficients for human–leopard and human–tiger conflict across background-sampling methods.** Standardized coefficients from full logistic regression models of (A) human–leopard conflict (HLC) and (B) human–tiger conflict (HTC) across the three background sampling methods at the optimal spatial scale and spatial cross validation block size. Estimates are presented as standardized log-odds coefficients with 95% confidence intervals. Coefficient sign indicates the direction of association with recorded conflict relative to background locations. The dashed vertical line represents the null association (β = 0). Continuous predictors were standardized prior to model fitting, such that coefficients represent the change in log odds associated with a one-standard-deviation increase in each predictor, conditional on the other predictors in the model. Predictors are ordered separately within each species by their mean coefficient across the methods in which they were retained. Missing symbols indicate predictors absent from the corresponding model. Intercepts are omitted.
